# Diphthamide formation in Arabidopsis requires DPH1-interacting DPH2 for light and oxidative stress resistance

**DOI:** 10.1101/2024.09.16.613322

**Authors:** Hongliang Zhang, Nadežda Janina, Koray Ütkür, Thirishika Manivannan, Lei Zhang, Lizhen Wang, Christopher Grefen, Raffael Schaffrath, Ute Krämer

**Author notes:** Department of Plant Biology, Carnegie Institution for Science, Stanford, CA 94305, USA. Address for correspondence, Professor Dr. Ute Krämer, Molecular Genetics and Physiology of Plants Ruhr University Bochum Universitaetsstrasse 150, Box 44, ND3/30, 44801 Bochum Germany.

## Abstract

Diphthamide is a post-translationally modified histidine residue of eukaryotic TRANSLATION ELONGATION FACTOR 2 (eEF2) and the target of diphtheria toxin in human cells. In yeast and mammals, the 4Fe-4S cluster-containing proteins Dph1 and Dph2 catalyze the first biosynthetic step of diphthamide formation. Here we identify *Arabidopsis thaliana* DPH2 and show that it is required for diphthamide biosynthesis, localizes to the cytosol and interacts physically with AtDPH1. Arabidopsis *dph2* mutants form shorter primary roots and smaller rosettes than the wild type, similar to *dph1* mutants which we characterized previously. Additionally, increased ribosomal -1 frameshifting error rates and attenuated TARGET OF RAPAMYCIN (TOR) kinase activity in *dph2* mutants also phenocopy the *dph1* mutant. Beyond the known heavy-metal hypersensitivity and heat shock tolerance of *dph1*, we newly show here that both *dph1* and *dph2* mutants are hypersensitive to elevated light intensities and oxidative stress, and that wild-type Arabidopsis seedlings accumulate diphthamide-unmodified eEF2 under oxidative stress. Both mutants share the deregulation of 1,186 transcripts in numerous environmental and hormone responses. AtDPH1 and AtDPH2 do not complement the corresponding mutants of *Saccharomyces cerevisiae*. In summary, DPH2 and DPH1 interact to function inter-dependently in diphthamide formation, the maintenance of translational fidelity, wild-type growth rates and TOR kinase activation, and they contribute to mitigating damage from elevated light intensities and oxidative stress. Under oxidative stress, a dose-dependent loss of diphthamide could potentiate downstream effects in a feed-forward loop. This work advances our understanding of translation and its interactions with growth regulation and stress responses in plants.

## Introduction

The eukaryotic translation elongation factor 2 (eEF2) mediates the translocation of the nascent peptidyl-tRNA from ribosomal A site to the P site during mRNA translation and protein biosynthesis (Gomez-Lorenzo et al., 2000). The eEF2 proteins of mammals, yeast, some archaea and Arabidopsis carry a unique post-translational modification on a conserved histidine residue named diphthamide (Su et al., 2013; Schaffrath et al., 2014; Zhang et al., 2022). Diphthamide was originally identified as the target of diphtheria toxin (DT), which catalyzes its ADP-ribosylation, resulting in the irreversible inactivation of eEF2, arrest of protein synthesis, and cell death (Honjo et al., 1968; Su et al., 2013; Schaffrath et al., 2014). The evolutionarily conserved multi-step pathway of diphthamide biosynthesis requires eight proteins, Dph1 to Dph8, in yeast, and at least seven proteins (Dph1 to Dph7) in mice and human (Honjo et al., 1968; Su et al., 2013; Schaffrath et al., 2014; Arend et al., 2023) (Fig. 1, Supplementary Fig. S1A). The first committed step of diphthamide biosynthesis is the transfer of a 3-amino-3-carboxypropyl (ACP) group from S-adenosylmethionine (SAM) onto the imidazole-C_2_ atom of a conserved His residue of eEF2 through an unconventional radical SAM reaction (Zhang et al., 2010; Dong et al., 2018; Fenwick et al., 2019) (Fig. 1A). In eukaryotes, this step is catalyzed by a complex of four Dph proteins, Dph1 to Dph4, possibly together with Dph8 (Liu et al., 2004; Webb et al., 2008; Zhang et al., 2010; Zhu et al., 2011; Thakur et al., 2012; Abdel-Fattah et al., 2013; Dong et al., 2014; Dong et al., 2017; Dong et al., 2018; Dong et al., 2019). Dph5 catalyzes the second step, the trimethylation of the amino group and mono-methylation of the carboxyl group of the ACP intermediate to generate diphthine methyl ester (Lin et al., 2014) (Supplementary Fig. S1A). Next, Dph7 mediates the de-methylation of the carboxyl group to produce diphthine (Su et al., 2012; Lin et al., 2014) (Supplementary Fig. S1A). Finally, the amidation of the carboxyl group is catalyzed by Dph6 to generate diphthamide (Su et al., 2012; Uthman et al., 2013) (Supplementary Fig. S1A). Despite the conserved, complex pathway of diphthamide biosynthesis, the function of diphthamide in cellular physiology is only partly elucidated. Several reports suggested that diphthamide plays a role in maintaining translational fidelity in yeast (Ortiz et al., 2006; Hawer et al., 2018) and mouse (Liu et al., 2012) by preventing -1 frameshift errors (Murray et al., 2016; Pellegrino et al., 2018). The lack of the diphthamide modification on eEF2 leads to severe growth and developmental defects in mice (Chen and Behringer, 2004) and humans (Alazami et al., 2015; Loucks et al., 2015; Hawer et al., 2020).

**Figure 1.**
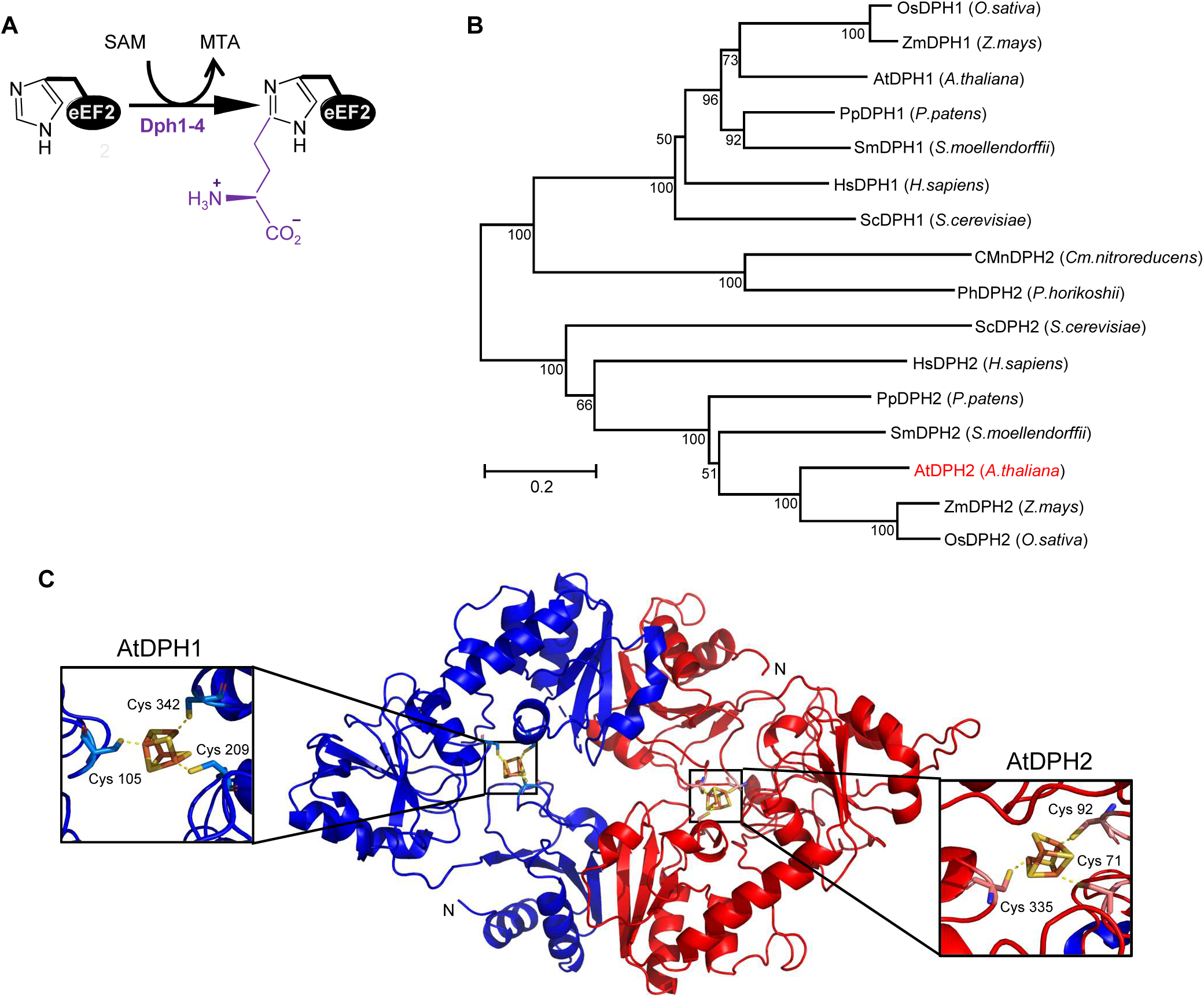
Evolutionary relationships of DPH1 and DPH2 homologs and predicted structure of the AtDPH1-AtDPH2 heterodimer. **A)** Scheme of first step of diphthamide biosynthesis (based on information from yeast and human). SAM: *S*-adenosylmethionine, MTA: 5’-methylthioadenosine. **B**) Neighbor-joining tree of DPH1 and DPH2 homologues. Numbers at the branching points indicate percent support from bootstrap analysis (1,000 iterations). Scale bar represents 0.2 substitutions per amino acid position (sum of branch lengths 6.10). **C**) Structural modeling-based prediction of the AtDPH1-AtDPH2 heterocomplex in Arabidopsis. The conserved Cys residues predicted to bind 4Fe-4S clusters are shown using a stick model (AtDPH1: Cys105, Cys209, and Cys342; AtDPH2: Cys71, Cys92, and Cys335). N marks the start of the amino acid sequences.

Only little is known about diphthamide in plants. We previously identified Arabidopsis homologs of yeast and human Dph1 to Dph7 by BLAST searches, supporting that the conserved diphthamide biosynthesis pathway may also exist in plants (Zhang and Krämer, 2018; Zhang et al., 2022) (Supplementary Fig. S1A). We further demonstrated that the diphthamide modification of Arabidopsis eEF2 (AT1G56070, also LOW EXPRESSION OF OSMOTICALLY RESPONSIVE GENES 1, LOS1) is indeed conserved in plants and requires AtDPH1 (Zhang et al., 2022). According to estimates based on tandem mass tag-based quantification, eEF2 ranks as the third most abundant protein after ATP-dependent Ribulose Bisphosphate Carboxylase (RuBisCo) ACTIVASE (RCA, AT2G39730) and the RuBisCo SMALL SUBUNIT 3B (RBCS-3B/S3B, AT5G38410) in the youngest leaf (LF 12) and the seventh most abundant protein in the youngest fully grown leaf (LF 7) of 22-d-old Arabidopsis (Col-0) plants (Mergner et al., 2020). Based on immunoblots and mass spectrometry, we concluded that approximately 96% of the eEF2 protein in Arabidopsis seedlings carries a diphthamide modification of H700 (Zhang et al., 2022). We reported that Arabidopsis *dph1* mutants lacking diphthamide show elevated translational -1 frameshifting rates and decreased growth compared to wild-type plants. Thus, the function of diphthamide in cellular biochemistry, as far as it is known in yeast and mammalian systems, is broadly conserved in Arabidopsis (Zhang et al., 2022).

Presently, the roles of other Arabidopsis proteins hypothesized to act in diphthamide biosynthesis remain unknown. In other eukaryotes, the non-canonical radical SAM reaction of the first step of diphthamide biosynthesis is catalyzed by Dph1-Dph4 (Zhang et al., 2010). Crystal structures of archaeal Dph2 of *Pyrococcus horikoshii* (PhDph2) and *Candidatus methanoperedens nitroreducens* (CmnDph2) revealed a homodimer binding one 4Fe-4S cluster per subunit (Zhang et al., 2010; Dong et al., 2018). Archaeal genomes do not encode Dph3 or Dph4 homologs, and PhDph2 alone can accomplish the initial step of diphthamide synthesis *in vitro* (Zhang et al., 2010). In eukaryotes, the 4Fe-4S cluster-containing Dph1 and Dph2 proteins are paralogues of the archaeal Dph2 protein, and they form a heterodimer in yeast (Dong et al., 2014). In the presence of dithionite as a reductant *in vitro*, the Dph1-Dph2 complex of yeast is sufficient for the first step of diphthamide biosynthesis (Dong et al., 2014). *In vivo*, Dph3 functions as an electron carrier for reducing the 4Fe-4S clusters of Dph1-Dph2 (Dong et al., 2014), and Dph4 serves as a putative J-type chaperone (Liu et al., 2004). Dph3 and Dph4, probably assisted by Dph8, are considered to chaperone Dph1-Dph2 for maintaining the 4Fe-4S cluster intact and in redox states required to form the ACP-modified EF2 in the first step (Schaffrath et al., 2014). It was demonstrated that in the archaeal Dph2 homodimer, only one 4Fe-4S cluster is necessary for activity in vitro (Zhu et al., 2011). Different from archaea, experiments in yeast revealed that the 4Fe-4S clusters of both the Dph1 and the Dph2 subunit are necessary for diphthamide biosynthesis *in vivo* (Dong et al., 2019). The cluster in Dph1 had catalytic activity, whereas the cluster in Dph2 was concluded to function in the reduction of the catalytic Dph1 cluster (Dong et al., 2019). In agreement with this, modeling predicted a conserved SAM binding site required for activity in all eukaryotic Dph1 proteins, which is absent in eukaryotic Dph2 proteins (Ütkur et al., 2023).

Here we characterize the function of a putative Dph2 orthologue in Arabidopsis (AT3G59630) employing two *dph2* T-DNA insertion mutants. We provide evidence supporting that AtDPH1 and AtDPH2 function as a heterodimer in the cytosol of Arabidopsis. The reduced-growth phenotype of Arabidopsis *dph2* knock-out mutants is similar to that of *dph1* mutants and of *dph1dph2* double mutants, thus indicating against any functional redundancy between AtDPH1 and AtDPH2. We newly report here that both Arabidopsis *dph1* and *dph2* mutants are hypersensitive to oxidative stress and elevated light intensities, and that diphthamide-unmodified eEF2 accumulates in wild-type Arabidopsis plants exposed to oxidative stress. Taken together, these results expand our knowledge on the biosynthesis and the biological roles of the diphthamide modification of eEF2, as well as its interactions with abiotic stress, in plants.

## Results

### DPH2 is conserved across eukaryotic species including land plants, and Arabidopsis *DPH2* and *DPH1* share similar expression patterns

In addition to demonstrating that AtDPH1 is required for diphthamide biosynthesis in the model plant *Arabidopsis thaliana*, we identified AtDPH2 to AtDPH8 as proteins hypothesized to contribute to diphthamide biosynthesis based on sequence similarity (Fig. S1A; Zhang et al., 2022). The amino acid sequence of AtDPH2 (AT3G59630) has around 25% identity with that of ScDPH2 and HsDPH2 (Supplementary Fig. S1A and B). According to a multiple sequence alignment, the three cysteine residues considered to be involved in 4Fe-4S cluster binding by known human and yeast DPH2 proteins are fully conserved in hypothesized DPH2 proteins from the bryophyte *Physcomitrium* (*Physcomitrella*) *patens*, the lycophyte *Selaginella moellendorffii*, rice and Arabidopsis (Supplementary Fig. S1B). A neighbor-joining tree supported a close relationship among putative DPH2 proteins from various eukaryotes, and it discriminated these from the DPH1 proteins present in all of these organisms (Fig. 1B). Interestingly, the single DPH1/2-related protein of archaea, PhDPH2 and CmnDPH2, grouped closer to the DPH1 orthologues (Fig. 1B). We modeled the predicted structure of a hypothesized heterodimeric AtDPH1-AtDPH2 complex using the known structure of the archaeal PhDPH2 (pdb/3LZD) as a template by homology modeling with default parameters (Bordoli et al., 2009; Waterhouse et al., 2018). The obtained model was consistent with the formation of a heterodimer carrying two 4Fe-4S clusters coordinated by conserved cysteine residues, as expected (AtDPH1: C105, C209, C342; AtDPH2: C71, C92, C335, Fig. 1C). The predicted AtDPH1-AtDPH2 heterodimer is composed of two protomers adopting approximately V shapes joined at the end of both arms, similar to the archaeal homodimer (Zhang et al., 2010). Several of the amino acids of AtDPH2 that are predicted to be positioned at the interface between protomers are conserved among known and putative eukaryotic DPH2 proteins, but not in AtDPH1 (Supplementary Fig. S1B; Supplementary Fig. S2).

### AtDPH2 physically associates with AtDPH1 in Arabidopsis

To test whether AtDPH2 can interact with AtDPH1, we performed Co-immunoprecipitation (Co-IP) and ratiometric bimolecular fluorescence complementation (rBiFC) assays (Grefen and Blatt, 2012; Xing et al., 2016). In Co-IP analyses upon transient expression in *Nicotiana benthamiana* leaf epidermal cells, AtDPH1 physically associated with AtDPH2, but not with AtDPH1 itself (Fig. 2A,B), consistent with the formation of a complex composed of AtDPH1 and AtDPH2 *in vivo*. The ability of AtDPH1 and AtDPH2 to interact with one another in plant cells was confirmed by rBiFC (Grefen and Blatt, 2012). We transiently co-expressed AtDPH1-nYFP with AtDPH2-cYFP or AtDPH1-cYFP in *N. benthamiana* leaf epidermal cells (Fig. 2C). Complementation of the YFP fluorescence was observed when AtDPH1-nYFP was co-expressed with AtDPH2-cYFP, but not with AtDPH1-cYFP, further supporting that AtDPH1 can physically interact with AtDPH2 (Fig. 2D,E). Together, these results suggest that AtDPH2 and AtDPH1 function as a heterodimeric complex.

**Figure 2.**
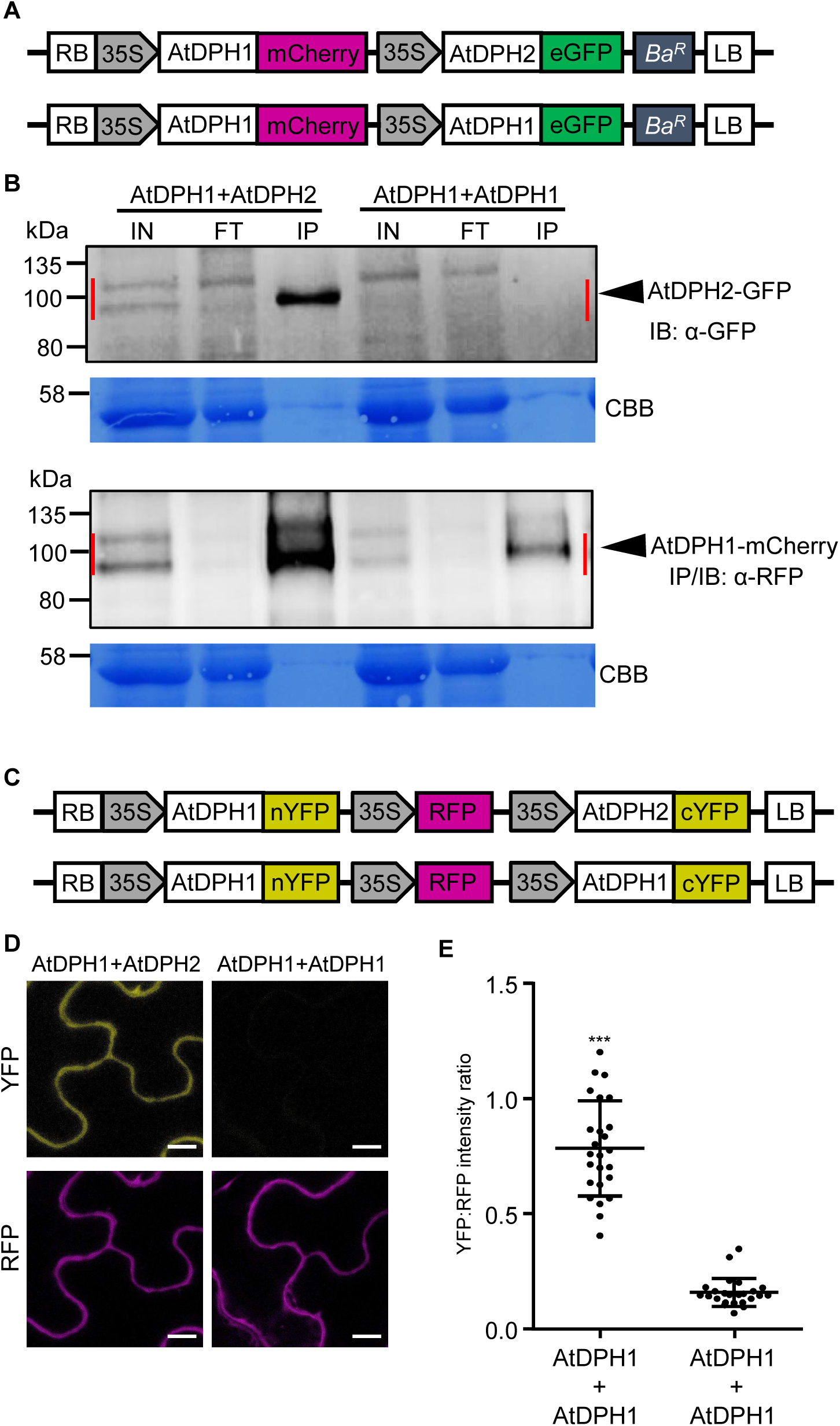
AtDPH1 interacts with AtDPH2 *in vivo*. **A)** Schematic representation of the 2in1 co-immunoprecipitation (Co-IP) constructs. **B)** Co-IP of AtDPH1 with AtDPH2 in transiently transfected *N. benthamiana* leaf epidermal cells (see **A**). Immunoprecipitation was carried out on protein extracts using anti-RFP beads, and protein interaction was detected by immunoblotting using an anti-GFP antibody. Blots stained with Coomassie Brilliant Blue (CBB) are shown below as loading controls. IN, input (total protein extract); FT, flow-through; IP, immunoprecipitate; IB: immunoblot. **C)** Schematic representation of the 2in1 ratiometric bimolecular fluorescence complementation (rBiFC) constructs. **D)** Representative confocal fluorescence microscopic images showing rBiFC analysis of AtDPH1 with AtDPH2 in transiently transfected *N. benthamiana* leaf epidermal cells (see **C**). Scale bars, 10 μm. **E)** Ratios of yellow:red fluorescence intensity (see **D**). Shown are mean ± s.d. (*n* = 25 different frames acquired from at least 3 independently infiltrated leaves in total). Total fluorescence intensities were measured by integrating over each frame in both the YFP and the RFP channels, subsequently ratioed, and ratios were plotted as a quantitative measure of YFP complementation. ***, *P* < 0.001 (two-tailed Student’s *t*-test compared with AtDPH1+AtDPH1 employed as a negative control).

### AtDPH2 is required for maintaining Arabidopsis growth

To characterize the functions of *AtDPH2*, we obtained two lines carrying T-DNA insertions in the *DPH2* locus, *dph2-1* and *dph2-2* (Supplementary Fig. S3A). We confirmed the genomic positions of the T-DNA insertions in the eighth exon out of ten exons in *dph2-1*, and in the predicted promoter in *dph2-2*, at the *DPH2* locus. RT-PCR and RT-qPCR revealed a lack of *DPH2* transcripts in *dph2-1* and reduced *DPH2* transcript levels in *dph2-2* (Supplementary Fig. S3B,C), suggesting that *dph2-1* is a knock-out mutant and *dph2-2* is a knock-down mutant of *DPH2*. For the genetic complementation of *dph2-1*, we generated stable homozygous *dph2-1* transgenic lines using a construct comprising a *DPH2* promoter fragment (1,800 bp upstream of the translational start codon) and the *DPH2* coding sequence translationally fused to a C-terminal *eGFP* coding sequence (*dph2-1 pDPH2:DPH2-GFP*). *AtDPH2* transcript levels in three independent *dph2-1 pDPH2:DPH2-GFP* complementation lines were similar to those in the wild type (Supplementary Fig. S3C). We additionally confirmed *DPH2* expression in the complemented lines by detecting the DPH2-GFP fusion protein with an anti-GFP antibody (Supplementary Fig. S3D).

In two-week-old Arabidopsis *dph1-1*, *dph2-1*, and double mutant *dph1 dph2* seedlings fresh biomass and primary root lengths were approximately 50% and 35% lower, respectively, than in the wild type and the *dph2-1 pDPH2:DPH2-GFP* complemented lines (Fig. 3A-C). Fresh biomass and primary root lengths of *dph2-2* were reduced by only 20% and 17%, respectively, compared to the wild type and the *dph2-1 pDPH2:DPH2-GFP* complemented lines (Fig. 3A-C), which was consistent with our observation of some residual *DPH2* transcript in the *dph2-2* mutant (see Supplementary Fig. S3B). Similar to seedlings grown on agar-solidified 0.5x MS medium on petri plates, soil-grown *dph2* mutants were smaller than the wild type and the complemented lines. Rosette sizes of 31-d-old soil-grown *dph2-1* and *dph1 dph2* were comparable to those of *dph1-1* (Supplementary Fig. S4A). These results suggest that *DPH2* is necessary for the maintenance of normal growth of Arabidopsis. The comparable extents of growth impairment in *dph1 dph2*, *dph1-1*, and *dph2-1* are consistent with our hypothesis that *DPH1* and *DPH2* operate in the same pathway. The aminoglycoside toxin hygromycin B inhibits polypeptide synthesis of bacteria, fungi and multi-cellular eukaryotes through binding to the nascent peptidyl-tRNA and preventing its eEF2-mediated translocation (Gonzalez et al., 1978). As is already known for yeast *dph1* to *dph7* mutants and the Arabidopsis *dph1* mutant, Arabidopsis *dph2-1* and *dph2-2* mutant seedlings were hypersensitive to hygromycin B (Supplementary Fig. S4B) (Hawer et al., 2018; Zhang et al., 2022).

**Figure 3.**
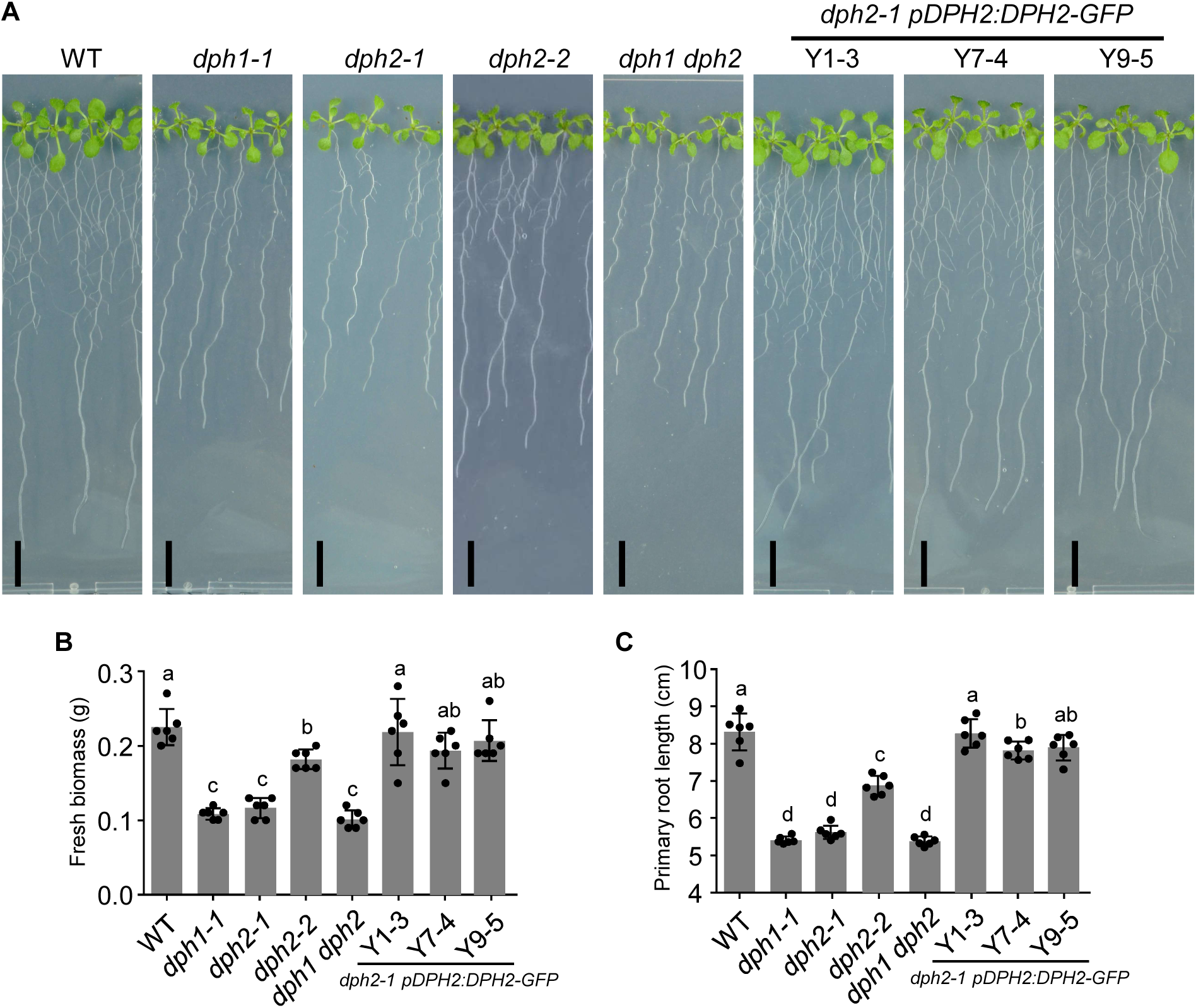
Smaller size of *dph1*, *dph2* and *dph1 dph2* double mutant seedlings. **A)** Photographs showing 2-w-old seedlings of two *dph2* mutant lines and of the *dph1-1 dph2-1* double mutant, alongside the wild type (WT) and the *dph1-1* mutant included as controls. Seedlings were cultivated in vertically oriented petri plates on agar-solidified 0.5x MS medium supplemented with 1% (w/v) sucrose. Scale bars, 1 cm. **B) C)** Bargraphs showing fresh biomass (**B**) and primary root length (**C**) of seedlings as shown in (**A**). Shown are mean ± s.d. (*n* = 6 replicate pools of 15 seedlings, with each pool cultivated on a replicate petri plate). Different characters represent significant differences between means (one-way ANOVA, Waller-Duncan test, *P* < 0.05).

### *AtDPH2* is ubiquitously expressed and its gene product localizes to the cytosol

Histochemical staining for β-glucuronidase (GUS) activity in *AtDPH2*-promoter (*pDPH2:GUS*) reporter lines suggested that the *AtDPH2* promoter is active in all the examined tissues, with elevated intensity of staining in meristematic regions of shoot and root tissues (Fig. 4A-G). Compared to the expression levels of *ACTIN8*, low levels of *AtDPH2* transcripts were detected in rosette leaf, cauline leaf, inflorescence, stem, root, and developing silique tissues (Fig. 4H). The ubiquitous expression pattern of *AtDPH2* is similar to that of *AtDPH1* (Zhang et al., 2022). To investigate the subcellular localization of AtDPH2, we imaged roots of *dph2-1 pDPH2:DPH2-GFP* lines by confocal microscopy. In root tips, the GFP signal was observed in the cytosol (Fig. 4I). In an independent approach, we generated a *pUBQ10:DPH2-GFP-T35S* construct and transfected mesophyll protoplasts of Arabidopsis wild-type plants for transient expression. Confocal microscopy revealed the presence of AtDPH2-GFP protein exclusively in the cytosol, while no fluorescence was observed in untransfected protoplasts (Supplementary Fig. S5). Together, these results show AtDPH2 is localized in the cytosol of most cells in the plant, alongside AtDPH1 (Zhang et al., 2022).

**Figure 4.**
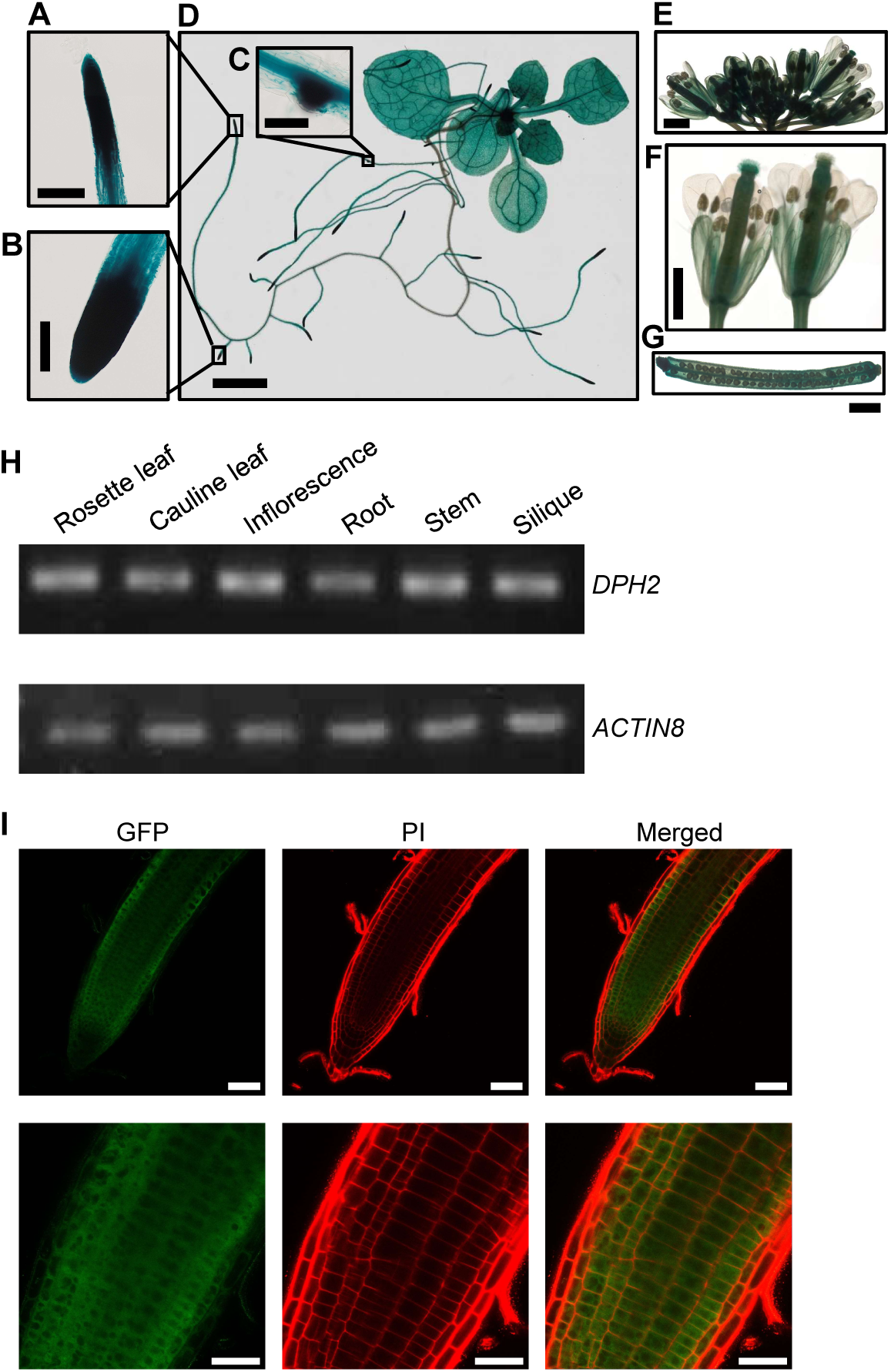
Expression pattern of *DPH2*, and DPH2 subcellular localization. **A**-**G)** Images showing histochemically detected β-glucuronidase (GUS) activity *pDPH2:GUS* transgenic plants (homozygous T3 generation, representative of 4 independently transformed lines). Insets show root tips at a higher magnification (**A-C**) of a 2-w-old seedling (**D**). Shoot apex (reproductive stage, **E**), flowers (**F**), and developing silique (**G**). Seedlings were grown on agar-solidified 0.5x MS medium containing 1% (w/v) sucrose (**A**-**D**), and all other tissues were harvested from 8-w-old soil-grown plants (**E**-**G**). Scale bars: 200 μm (**A**), 50 μm (**B**), 2 mm (**C**, **G**), 100 μm (**D**), 1 mm (**E**, **F**). **H)** Detection of *DPH2* transcript by RT-PCR (30 cycles) in different tissues of 7-w-old soil-grown Arabidopsis (Col-0). *ACTIN8* was used as a housekeeping control gene (26 cycles). **I)** Representative confocal laser scanning microscopic images of a root tip of a 9-d-old *dph2-1 pDPH2:DPH2-GFP* (line Y1-3) seedling stained with propidium iodide (PI, red). Scale bars, 50 μm (upper row), 25 μm (lower row).

### AtDPH2 is necessary for diphthamide biosynthesis and for maintaining wild-type levels of TARGET OF RAPAMYCIN (TOR) kinase activity

Specific antibodies are available to detect either diphthamide-unmodified eEF2 or global eEF2 protein, comprising both the unmodified and the modified form of eEF2, in protein extracts from human and yeast cells (Stahl et al., 2015; Hawer et al., 2018). We previously demonstrated that these antibodies can be used to detect the corresponding forms of Arabidopsis eEF2 (Zhang et al., 2022). As was shown previously for *dph1* using immunoblotting, diphthamide-unmodified eEF2 was also detected in *dph2* single and *dph1 dph2* double mutants, but not in the wild type and in the *dph2-1 pDPH2:DPH2-GFP* complemented lines (Fig. 5A). According to band intensities, the amount of unmodified eEF2 detected in *dph2-2* was approximately 70% of the amount of unmodified eEF2 observed in *dph1-1*, *dph2-1*, and *dph1 dph2* mutants (Fig. 5A), suggesting that part of the eEF2 pool carries the diphthamide modification in the *dph2-2* mutant. To examine whether, in analogy to DPH1, DPH2 is also required for maintaining translational fidelity in Arabidopsis, we measured ribosomal -1 frameshifting rates using the -1 frameshifting dual-luciferase reporter developed earlier (Zhang et al., 2022). We transiently transfected aliquots of mesophyll protoplasts of the wild-type, *dph2* single, and *dph1 dph2* double mutants either with a control reporter for in-frame translation or a test reporter for programmed -1 frameshifting (Fig. 5B) (Zhang et al., 2022). Compared to the wild type, -1 frameshift error rates were elevated by around 57% on average in both the *dph2-1* single and *dph1 dph2* double knock-out mutants, and by 29% in the knock-down mutant *dph2-2* (Fig. 5C; Supplementary Fig. S6D).

**Figure 5.**
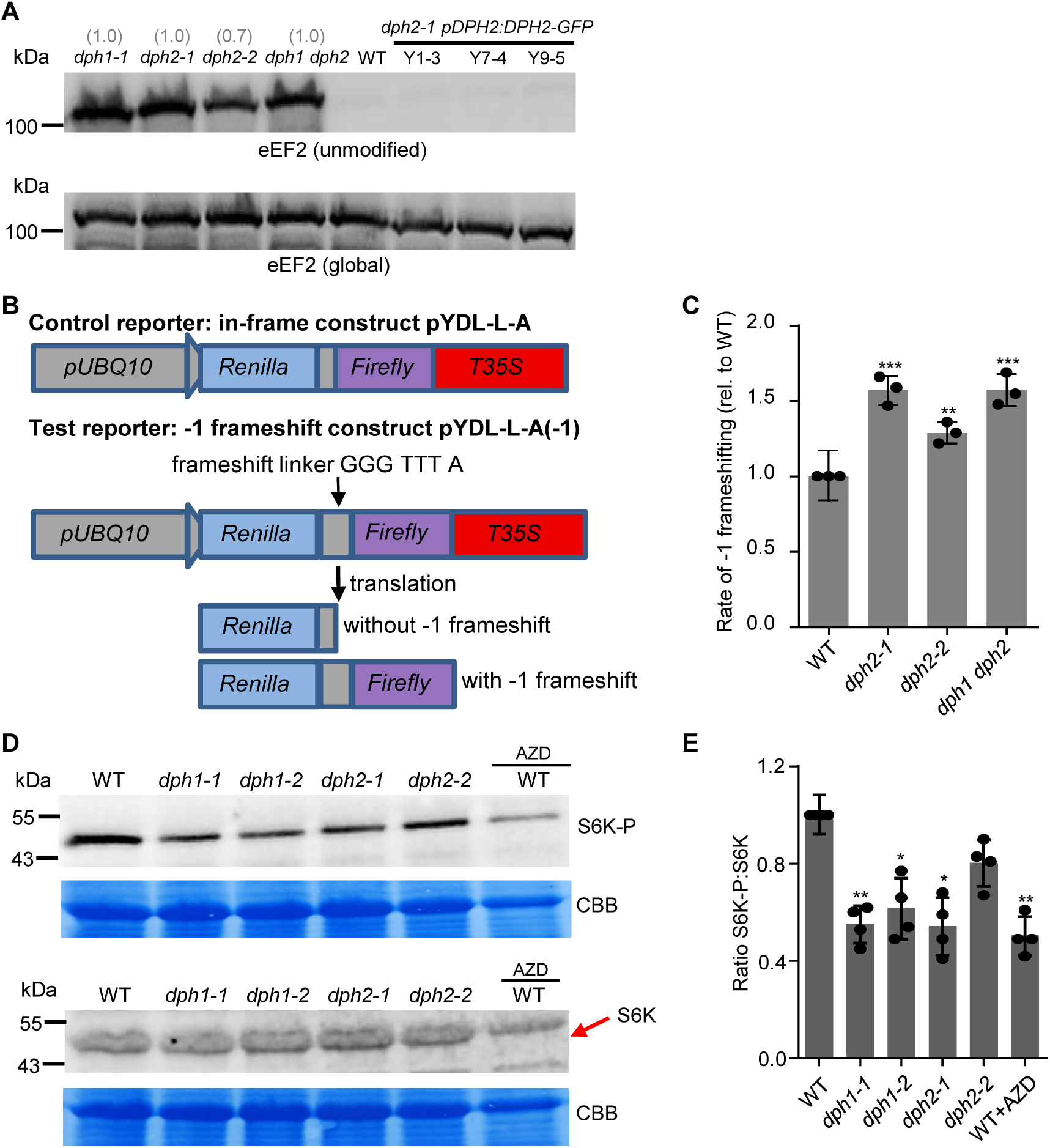
Diphthamide modification of eEF2, translational reading-frame accuracy, and TOR kinase activity depend on both AtDPH1 and AtDPH2. **A)** Immunoblot probed with an antibody specific for detecting eEF2 lacking the diphthamide modification (unmodified eEF2), and an antibody recognizing all forms of eEF2 (global eEF2). Total protein extracts from equal amounts of shoot biomass of 14-d-old seedlings grown in agar-solidified 0.5x MS medium containing 1% (w/v) sucrose were resolved by SDS-PAGE and blotted. Numbers above the upper blot image show band intensities for unmodified eEF2 in a given lane, relative to that for *dph1-1*. **B)** Schematic representation of the -1 frameshifting error reporter system, which consisted of pYDL-L-A(-1) as a test reporter and pYDL-L-A as a control reporter. Both reporters encode a chimeric fusion protein of renilla luciferase and firefly luciferase, separated by a linker. For the control reporter, the firefly luciferase is in frame with renilla luciferase. For the test reporter, the firefly luciferase is in -1 frame relative to renilla luciferase so that a fusion protein is only produced if -1 frameshifting occurs during the translation of the linker sequence. **C)** Boxplot showing the frequency of -1 frameshifting during translation of the test reporter transcript for each genotype relative to the wild type (see **B**). To obtain -1 frameshifting error rates, we divided the ratio of firefly:renilla luciferase activity of the test -1 frameshift reporter construct by that of the in-frame control reporter construct for each genotype (see Fig. S6A). Luciferase activities were quantified in transiently transfected Arabidopsis leaf mesophyll protoplasts prepared from 3-week-old soil-grown *dph2* mutant, *dph1 dph2* double mutant and wild-type (WT) plants. Shown are mean ± s.d. (*n* = 3 independently transfected replicate aliquots of protoplasts). Significant differences from WT: **, *P* < 0.01, ***, *P* < 0.001 (one-way ANOVA with Tukey’s test). **D**-**E)** Reduced TOR activity in *dph1* and *dph2* mutants. Representative immunoblots (**D),** and bargraph of TOR kinase activity as reflected by ratios of S6K-P:S6K band intensities (**E**). Total protein was extracted from equal amounts of 14-d-old seedlings grown in liquid 0.5x MS medium supplemented with 0.5% (w/v) sucrose, resolved by SDS-PAGE, blotted and probed with antibodies against S6K-P and S6K, respectively. WT seedlings treated with 2 μM TOR inhibitor AZD-8055 (AZD) for 1 d were used as controls. The position of the S6K band is indicated by a red arrow (**D**). Data are mean ± s.d., *n* = 4 independent experiments (**E**, see also Fig. S6B-D). Significant differences from WT: *, *P* < 0.05, **, *P* < 0.01, one-way ANOVA with Games-Howell test.

Target of rapamycin (TOR) kinase (AT1G50030) is a central regulator controlling plant growth, development and responses to environmental stresses (Shi et al., 2018; Zhang et al., 2022). TOR activity was significantly decreased in 14-d-old *dph2* seedlings compared to wild-type seedlings, based on the ratio of phosphorylated ribosomal-protein S6 kinase S6K (S6K-P) to S6K protein (AT3G08720) amounts as a read-out (Fig. 5D and E; Supplementary Fig. S6A-C) (Busche et al., 2021). Ratios of S6K-P to S6K were reduced to approximately 55% of wild-type ratios in *dph1-1* and *dph1-2* as well as in *dph2-1*, comparable to the ratios observed in the wild type treated with the TOR kinase inhibitor AZD-8055 (Montané and Menand, 2013), and to about 80% of wild-type ratios in *dph2-2* (Fig. 5E) (see also Zhang et al., 2022). Taken together, these results demonstrate that diphthamide modification of eEF2 requires AtDPH2 in addition to AtDPH1, and that both of them are necessary for maintaining translational fidelity and TOR activity in Arabidopsis.

### Arabidopsis *dph1* and *dph2* mutants are hypersensitive to oxidative stress and elevated light intensity

Several studies suggested that diphthamide plays a role in the response to oxidative stress in humans (Argüelles et al., 2013; Argüelles et al., 2014). For example, *dph* knock-out mutants of MCF7 cells accumulated high levels of ROS under normal growth conditions, implying a disruption of cellular redox state (Mayer et al., 2019). In line with this, it was noted that diphthamide-less archaea and eukaryotes (parabasalids) are obligate anaerobes (Narrowe et al., 2018). In our previous transcriptomic analysis of Arabidopsis *dph1* mutants, the GO term “oxidative stress” was significantly enriched among transcripts up-regulated in two independent *dph1* mutant lines compared to the wild type (Zhang et al., 2022).

We tested the sensitivity of *dph1* and *dph2* mutants to oxidative stress. The herbicide methyl viologen (MV, paraquat) causes light-dependent oxidative stress in plants (Foyer et al., 1994), and it inhibited growth of the wild-type, *dph1-1*, and *dph2-1* seedlings, with reduced size and chlorotic leaves (Fig. 6A). Upon exposure to 12 µM MV for 3 d, the seedlings of the mutants were completely or almost completely bleached, whereas leaves of wild-type seedlings remained mostly green (Fig. 6A). In more detail, chlorophyll contents of *dph1-1* and *dph2-1* seedlings were lower than those of the wild type before the onset of MV treatment, and MV treatment reduced chlorophyll contents of all genotypes (Fig. 6B). Importantly, chlorophyll contents of MV-exposed *dph1-1* and *dph2-1* seedlings normalized to those of untreated controls were lower than observed for the wild type (Fig. 6C). These data suggest that *dph1-1* and *dph2-1* seedlings are more sensitive to MV treatment than wild-type seedlings.

**Figure 6.**
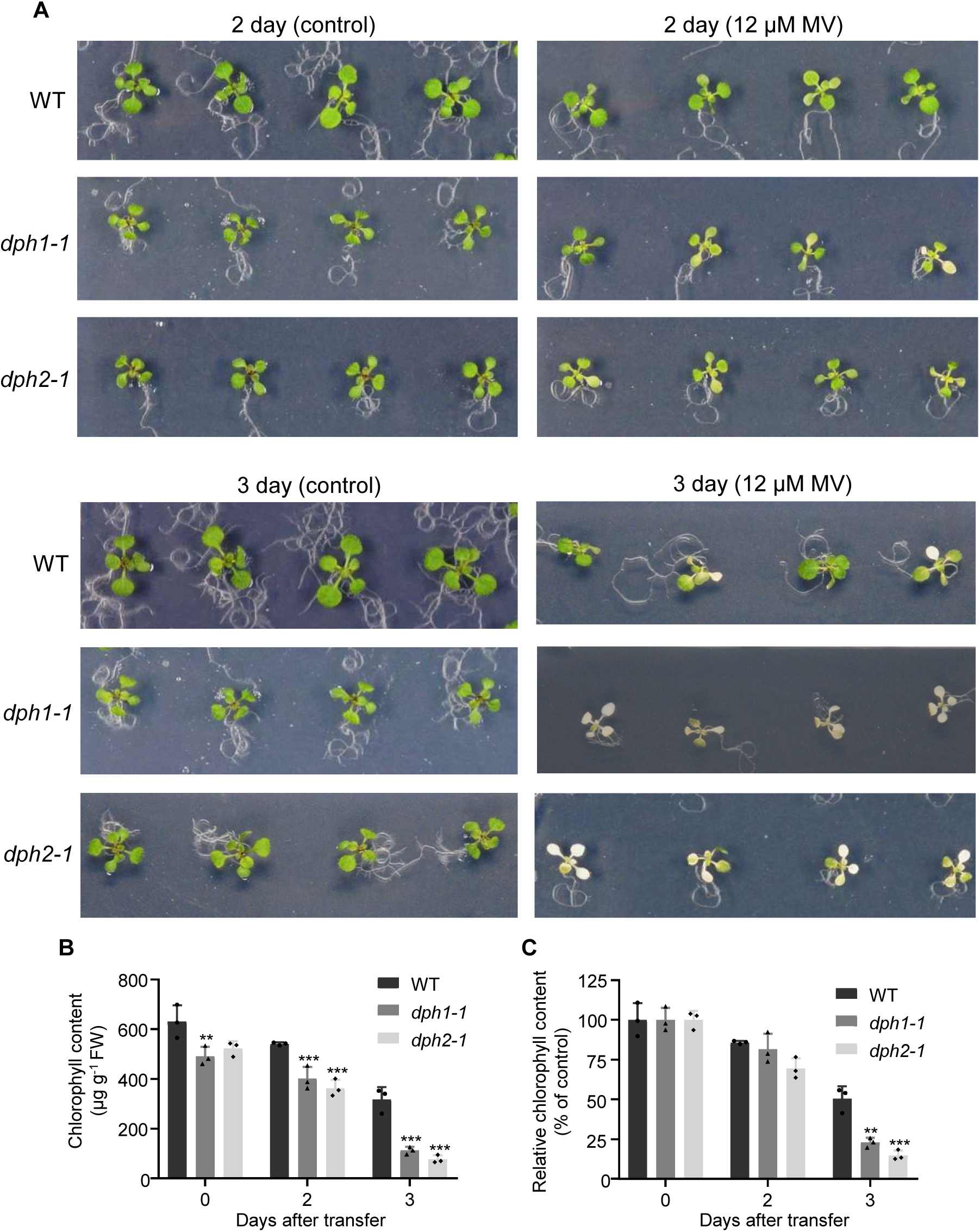
Methyl viologen sensitivity assay. **A)** Photographs of the wild-type (WT), *dph1*-*1*, and *dph2*-*1*. Seedlings were pre-cultivated on agar-solidified 0.5x MS medium supplemented with 1% (w/v) sucrose in vertical orientation for 11 d before transfer to fresh 0.5x MS medium or 0.5x MS supplemented with 12 µM methyl viologen (MV). Photos of 4 representative seedlings per genotype were taken after 2 d (upper panel) and 3 d (lower panel) of treatment in horizontal orientation. **B, C)** Chlorophyll content of the seedlings shown in (**A**). Data are mean ± s.d. (*n* = 3 petri plates, with tissues from 12 to 16 seedlings grown on one plate pooled per replicate). Relative chlorophyll contents (**C**) were calculated by normalizing to the mean of same genotype under the control condition (0 d). Significant differences from WT at the same time point: **, *P* < 0.01, ***, *P* < 0.001, based on one-way ANOVA with Tukey’s test (**B**, **C**). FW: fresh biomass.

In addition, we observed that *dph1-1*, *dph2-1* and *dph1 dph2* seedlings are hypersensitive to elevated light intensities (145 µmol photons m^-2^ s^-1^), reflected by smaller rosettes and a more chlorotic appearance than in the control light conditions (100 µmol photons m^-2^ s^-1^) employed by us or in wild-type seedlings cultivated under elevated-light conditions (Fig. 7A). Fresh biomass and chlorophyll contents of the wild type remained similar following cultivation under both our control and elevated-light illumination conditions. In contrast, upon cultivation of *dph1-1*, *dph2-1* and *dph1 dph2* mutant seedlings in elevated light, fresh biomass and chlorophyll contents were reduced to between 60% and 70% of those observed in the respective genotype grown under our control illumination conditions (Fig. 7B-E). When cultivated on soil under elevated light (185 µmol photons m^-2^ s^-1^, constant light) but not in control conditions (120 µmol photons m^-2^ s^-1^, 16 h/8 h day/night cycles), several leaves of *dph1-1*, *dph2-1*, and *dph1 dph2* mutants developed yellowing whereas the leaves of wild-type plants remained green (Supplementary Fig. S7). Taken together, these results show that *dph1* and *dph2* mutants are hypersensitive to elevated light intensities and to oxidative stress generated by MV treatment.

**Figure 7.**
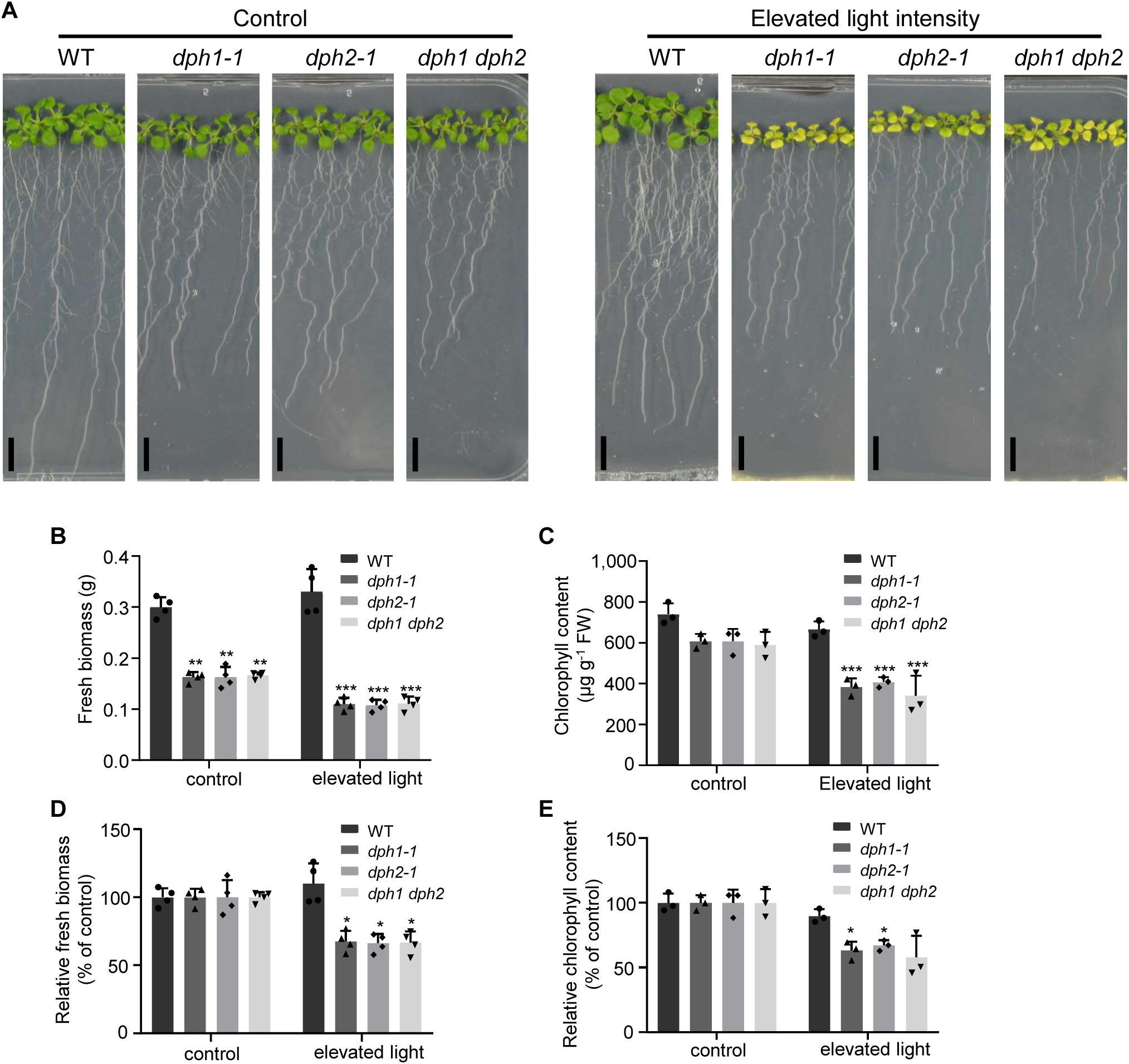
Test for sensitivity to elevated light intensity. **A)** Photographs of 14-d-old wild-type (WT), *dph1-1*, *dph2-1* and *dph1 dph2* mutants cultivated under control conditions (100 µmol photons m^-2^ s^-1^) and under elevated light intensity (145 µmol photons m^-2^ s^-1^) in long days. Seedlings of each genotype pre-grown on agar-solidified 0.5x MS medium with 1% (w/v) sucrose under control conditions were either maintained under the same conditions or transferred into to elevated light for 7 d, with cultivation in vertically oriented petri plates. Scale bars, 1 cm. **B)** Fresh biomass of the seedlings shown in (**A**). **C)** Chlorophyll content of the seedlings shown in (**A**). FW: fresh biomass. Data are mean ± s.d. (*n* = 4 petri plates for **B**; *n* = 3 petri plates for **C**), with tissues from 18 seedlings pooled per replicate plate. **D**-**E)** Relative fresh biomass (**D**) and chlorophyll content (**E**) of the seedlings under control or an elevated light intensity as shown in (**A**). Data shown are normalized to the mean of same genotype under the control condition. Significant differences from WT under each growth condition: *, *P* < 0.05, **, *P* < 0.01, ***, *P* < 0.001, based on one-way ANOVA with Games-Howell test (**B-E**) or Tukey’s test (**C**).

### Arabidopsis wild-type seedlings exposed to oxidative stress accumulate diphthamide**-**unmodified eEF2

Earlier, we observed that exposure to the heavy metals copper and cadmium caused a strong decrease in the diphthamide-modified pool of eEF2 in Arabidopsis (Zhang et al., 2022). Here we tested whether other abiotic stresses affect diphthamide biosynthesis, as well. Unmodified eEF2 accumulated in wild-type seedlings exposed to 12 µM MV for 2 d, but not in seedlings cultivated in elevated light (145 µmol photons m^-2^ s^-1^) for 1 w (Fig. 8A). To investigate the effect of MV in more detail, we cultivated wild-type seedlings on 0.5x MS medium supplemented with a range of MV concentrations for 2 d. Leaf chlorosis intensified (Fig. 8B), and shoot chlorophyll contents dropped (Fig. 8C), in a dose-dependent manner. The amount of unmodified eEF2 increased gradually with rising MV concentration in the medium, while the levels of global eEF2 began to decrease moderately at the higher MV concentrations tested (Fig. 8D). The relationship between chlorophyll contents and the ratios of unmodified-to-global eEF2 was approximately linear (Fig. 8E).

**Figure 8.**
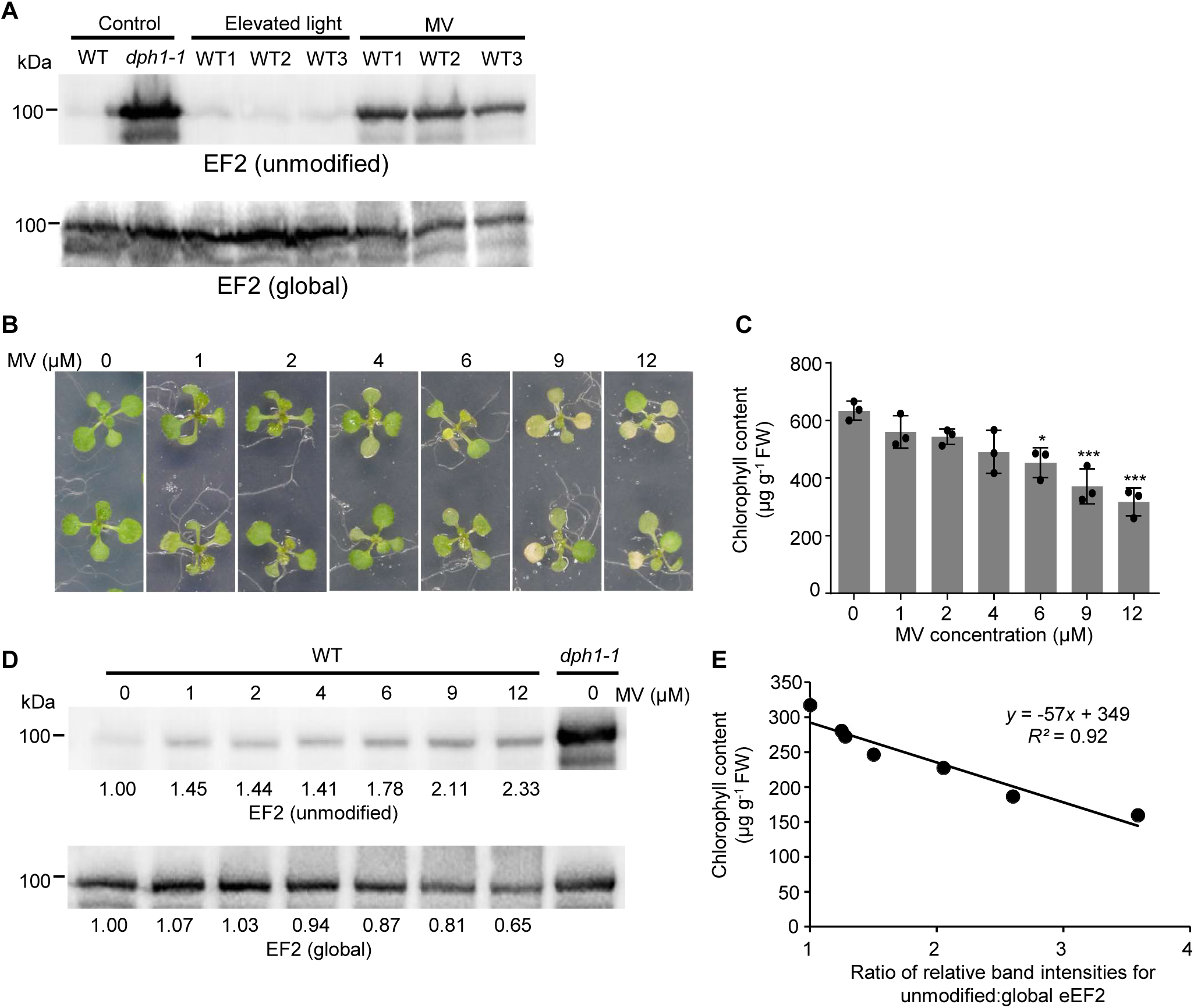
Diphthamide-unmodified eEF2 protein accumulates in wild-type seedlings exposed to methyl viologen. **A)** Immunoblot detection of unmodified and global eEF2 protein in total protein extracts from 14-d-old wild-type (WT) and *dph1-1* seedlings exposed to an elevated light intensity (see Fig. 7) and 12 µM MV (see **B**). Samples from stress treatments of 3 independent experiments were included (one lane per experiment). **B)** Photographs of representative WT seedlings after cultivation on medium containing a series of MV concentrations for 2 d. Twelve-d-old seedlings were transferred to the same medium supplemented with MV (see Fig. 6). **C)** Chlorophyll content of the shoots of seedlings shown in (**B**). Data are mean ± s.d. (*n* = 3 replicates, with 10 seedlings grown on one plate pooled per replicate). Significant differences from WT seedlings grown without MV: *, *P* < 0.05, ***, *P* < 0.001 (one-way ANOVA, Tukey’s test). **D)** Immunoblot detection of unmodified and global eEF2 protein in total protein extracts from seedlings shown in (**B**). *dph1-1* seedlings grown without MV treatment were included as a control. Band intensities were quantified by ImageJ. Numbers below blot images indicate band intensities normalized to those of WT seedlings grown on medium without MV. **E)** Plot showing the quantitative relationship between unmodified eEF2 protein and chlorophyll content in WT seedlings exposed to a range of MV concentrations (see **B**-**D**). The equation shown is based on linear regression analysis. FW: fresh biomass.

To investigate the way in which diphthamide biosynthesis was partially inhibited by oxidative stress, we quantified the transcript levels of *DPH1*, *DPH2*, and the other putative *DPH* genes (*DPH3* to *DPH7*), in wild-type seedlings by RT-PCR. In parallel, we analyzed protein levels of the DPH1-GFP and DPH2-GFP fusion proteins in *dph1-1* and *dph2-1* complemented lines by immunoblots. DPH1-GFP and DPH2-GFP protein levels were reduced by about one-third in seedlings exposed to 12 µM MV for 2 d, whereas *DPH* gene transcript levels remained largely unchanged (Supplementary Fig. S8A and B). Consequently, oxidative stress affected either the synthesis or the stability of the DPH1 and DPH2 proteins, and it likely affected their biochemical activity even more strongly (see Fig. 8E).

### Transcriptomic changes in *dph1-1* and *dph2-1* mutants

To examine the broader consequences of the lack of diphthamide, we conducted sequencing-based transcriptomics and compared the *dph1-1* and *dph2-1* mutants with wild-type seedlings in three independent experiments. Mutant growth is the most strongly impaired at the young seedling stage, and it gradually catches up with that of the wild type with increasing plant age. Consequently, here we sampled two-week-old seedlings, different from a previous study on rosette tissues of five-week-old soil-grown *dph1* mutant (*dph1-1*, *dph1-2*) and wild-type plants (Zhang et al., 2022).

Principal component analysis (PCA) separated the transcriptomes of wild-type seedlings from those of both the *dph1-1* and *dph2-1* mutant seedlings along the first PC explaining 80% of the variance (Fig. 9A). The second PC explained 9% of the variance and separated the *dph1-1* from the *dph2-1* samples. Average normalized transcript abundance of *DPH1* was around 10 transcripts per million (TPM) in both the wild type and the *dph2-1* mutant, and it was reduced to 0.6 TPM in the *dph1-1* mutant, below the threshold of 1.46 TPM used to filter out non-expressed genes (Supplemental Data Set 1, Fig. 9B). Similarly, *DPH2* was at 10 TPM in the wild type and in the *dph1-1* mutant, and down to 1.8 TPM in the *dph2-1* mutant (Supplemental Data Set 1, Fig. 9C). These data confirmed complete or near-complete loss-of-function in the respective mutant genotypes.

**Figure 9.**
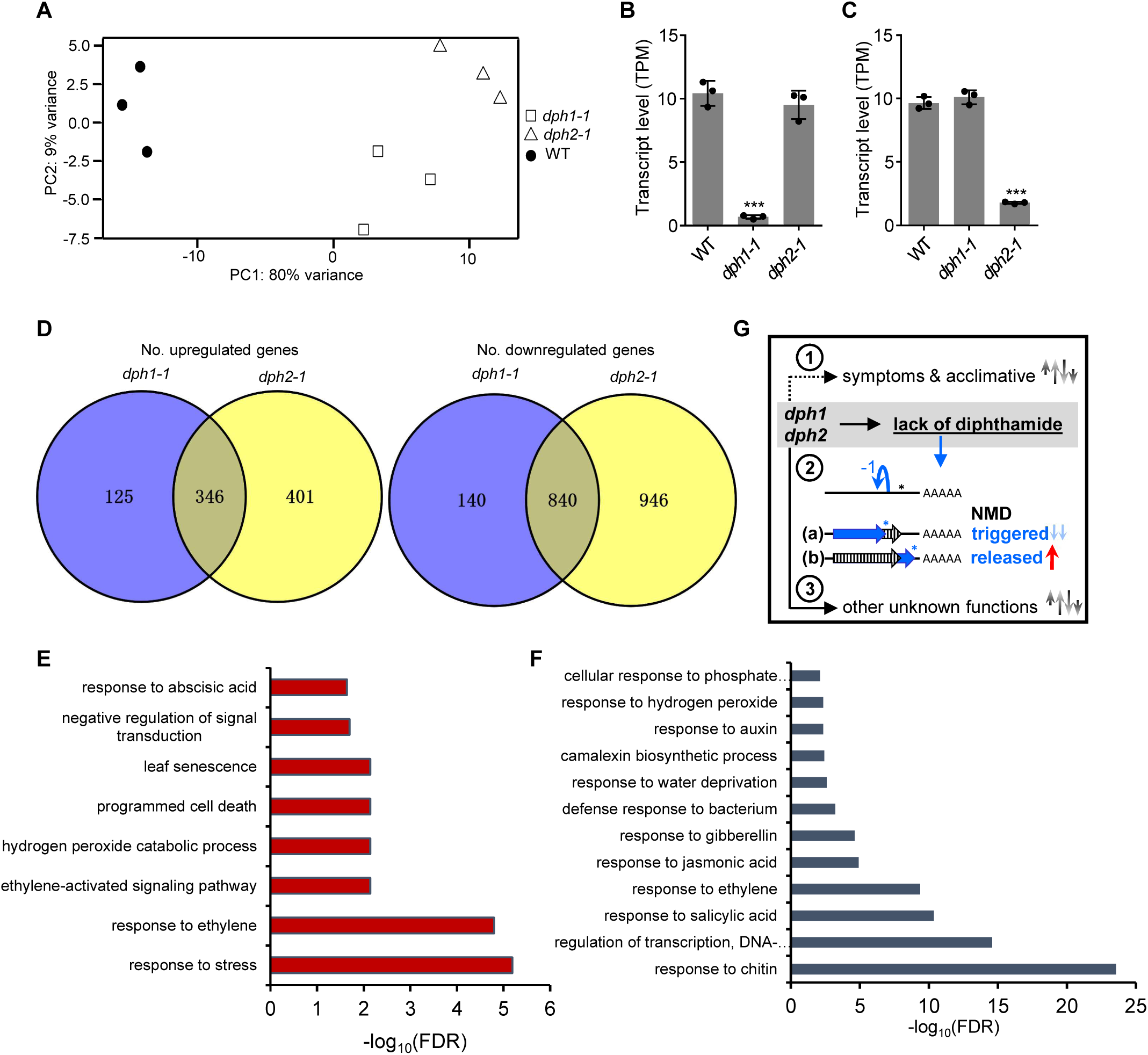
Transcriptome alterations in *dph1-1* and *dph2-1* mutants compared with wild-type seedlings. **A)** Principal Component Analysis of transcriptome sequencing data (based on transcripts per kilobase million, TPM). **B, C)** TPM of *DPH1* (**B**) and *DPH2* (**C**) in WT, *dph1-1*, and *dph2-1* mutants. Data are mean ± s.d. Significant differences from WT: ***, *P* < 0.001 (one-way ANOVA, Tukey’s test). **D**, Venn diagrams of transcripts present at higher (upregulated) and lower (downregulated) levels in *dph1-1* and *dph2-1* mutants compared to the wild type (WT). **E, F)** Representative GO terms significantly enriched among genes of which transcript levels were upregulated (**E**) and downregulated (**F**) in both *dph1-1* and *dph2-1* mutants compared to WT (FDR < 0.05). **G)** Model outlining hypotheses of alterations in the transcriptomes of the mutants, with indirect (1) and direct (2, blue) effects of the lack of diphthamide, as well as hypothetically possible other unknown functions of DPH1 and DPH2 (3). For the vertical arrows on the right (**G**), directions indicate up- or downregulation, the number of arrows symbolizes the number of genes undergoing, and the lengths of arrows the magnitude of, expected changes in transcript levels. Data shown are from *n* = 3 independent experiments, with samples of 2-week-old seedlings cultivated in liquid 0.5x MS medium supplemented with 0.5% (w/v) sucrose (**A**-**F**).

Transcript levels of 1,186 genes were differentially abundant in both the *dph1-1* and the *dph2-1* mutant when compared to the wild type, and of these, 346 were upregulated and 840 were downregulated in the mutants (Fig. 9D, Supplemental Data Set 2). Among the 346 transcripts that were more abundant in both the *dph1-1* and the *dph2-1* mutant than in the wild type, most of the significantly enriched biological processes are associated with abiotic stress responses, such as “response to ethylene and abscisic acid”, and “leaf senescence” (Fig. 9E, Supplemental Data Set 3). The GO terms hydrogen peroxide metabolism and catabolism (GO:0042744, GO:0042743), as well as reactive oxygen species (ROS) metabolism (GO:0072593), and response to oxidative stress (GO:0006979) were also significantly enriched among these transcripts (Supplemental Data Set 3), consistent with the increased sensitivity of *dph1* and *dph2* mutants to oxidative stress. Among the 840 transcripts identified here to be less abundant in both the *dph1-1* and the *dph2-1* mutant than in the wild type, biotic stress responses were significantly enriched, for example “response to chitin” (GO: 0010200), “response to salicylic acid” (GO: 0009751), and “response to ethylene” (GO: 0009723), as well as “regulation of DNA-templated transcription” (GO: 0006355; Fig. 9F, Supplemental Data Set 3). Besides these common differentially expressed genes, 14% of the upregulated and 7% of downregulated transcripts were unique to *dph1-1*, and much larger proportions of 46% and 49%, respectively, were unique to *dph2-1*.

It is important to consider that in *dph1* and *dph2* mutants there are several possible causes of transcriptome alterations. We expect that a large proportion of transcriptome differences between the mutants and the wild type arise indirectly either as a symptom of, or from the mutants’ attempts to compensate, the consequences of the absence of the diphthamide modification on eEF2 (marked as 1 in Fig. 9G). However, the lack of diphthamide can also have more direct effects on the transcriptome (marked as 2 in Fig. 9G). For a protein-coding transcript, ribosomal -1 frameshifting error can result in either an early termination of translation or a translational stop codon read-through, with opposing effects on the length of the 3’ UTR of the affected transcript. Because of a decreasing likelihood of stop codon read-through with increasing distance of the frameshift from the stop codon on a given transcript, early termination is generally the more likely event and can thus be predicted to affect a larger number of transcripts compared to the number of transcripts affected by stop codon read-through (2a in Fig. 9G). Transcripts harboring unusually long 3’ UTRs exceeding 350 nt undergo nonsense-mediated decay (NMD) unless protected by specific sequence characteristics (Kertész et al., 2006; Kalyna et al., 2012; May et al., 2018).

Although ribosomal -1 frameshifting error rates in our synthetic reporter construct of a single slippery site only were about one in three hundred (0.33%, see Fig. 5C) in *dph1* and *dph2* mutants, quantitatively large alterations in the levels of specific transcripts are possible. For instance, even if only a small proportion of the total amount of transcript of a specific gene were released from strong NMD occurring in the wild type, the level of this transcript could increase substantially in the mutants (conceivable for transcripts derived from pseudogenes containing -1 frameshift mutations, for example; 2b in Fig. 9G). By comparison to this, we expect smaller magnitudes of the downward changes in the abundances of transcripts when NMD is triggered by frameshifting in the mutants (2a in Fig. 9G).

For any plant-endogenous transcript undergoing higher -1 frameshifting rates in the mutants than we observed using our synthetic reporter construct, we expect an enlarged difference in abundance between the mutants and the wild type. Finally, we cannot exclude that the DPH1 or DPH2 protein themselves, have additional, yet unidentified, regulatory functions in Arabidopsis independent of the diphthamide modification of eEF2 (marked as 3, Fig. 9G).

Globally, across the total number of differentially abundant transcripts in both of the mutants, a moderate downregulation in the mutants predominated, as would be expected if NMD acted on an increased number of transcripts in the mutants (Fig. 9D, 2a in Fig. 9G). We observed the largest magnitudes of changes among transcripts upregulated in *dph1-1*, *dph1-2* (Zhang et al., 2022) and *dph2-1* when compared to the wild type: two transposable elements (full-length TE *Sadhu3-2*, AT3G42658; Ty1 COPIA-like retrotransposon AT5G35935; Supplemental Data Set 2). There were a few additional similar examples (6 COPIA-like retrotransposons, 2 Gypsy-like retrotransposons, for example), including also 8 potential natural antisense transcripts. Transcript level alterations for these genes conformed to expectations for transcripts undergoing NMD in the wild type that is partially released in the mutants (see 2b in Fig. 9G).

Transcript levels of *SMG7* (AT5G19400, *SUPPRESSOR WITH MORPHOGENETIC EFFECT ON GENITALIA 7*), which has a central role in NMD, were elevated in *dph1-1* and *dph2-1* compared to the wild type (Supplemental Data Set 2), consistent with a general activation of the NMD pathway (Raxwal and Riha, 2023). However, levels of the transcript encoding another central NMD protein, UPF1 (AT5G47010), were unaltered in the mutants, although Araport (https://araport.org/) suggests an overall high degree of co-expression with *SMG7*.

Among the transcripts exhibiting reduced levels in *dph1-1* and *dph2-1*, those encoding class II trehalose 6-phosphate synthase-like proteins of unknown function (TPS8, TPS9 and TPS11) are noteworthy (Supplemental Data Set 2), given their possible roles in metabolic regulation and the decrease in TOR kinase activity observed in *dph1* and *dph2* mutants (see Fig. 5D,E) (Fichtner and Lunn, 2021; Zhang et al., 2022). In addition, transcript levels of genes encoding regulators of the circadian clock and of light responses were downregulated in the mutants, for example *CCA1* (AT2G46830), *LHY* (AT1G01060), *PRR9* (AT2G46790), *CDF1* (AT5G62430), *PCC1* (AT3G22231), *RVE2* (AT5G37260), *LCL1* (AT5G02840), *TZP* (AT5G43630), *HY5* (AT5G11260), *SPA1* (AT2G46340)*, SPA3* (AT3G15354) and *SPA4* (AT1G53090) (Fittinghoff et al., 2006; McClung, 2019) (Supplemental Data Set 2).

Finally, transcript levels of several iron deficiency response genes were decreased in the mutants, including *bHLH038* (AT3G56970), *bHLH101* (AT5G04150), *IRT1* (AT4G19690, only in *dph2-1*), *FRO2* (AT1G01580). Transcript levels of Fe homeostasis genes *FRO6* (AT5G49730) and *FRO7* (AT5G49740) were also reduced, accompanied by an upregulation of Fe sufficiency marker genes *FER4* (AT2G40300) and *VTL2* (AT1G76800) (Supplemental Data Set 2). This de-regulation of Fe homeostasis-related transcripts was likely a consequence of the perception of an overall more Fe-sufficient physiological status in the *dph1* and *dph2* mutants (Krämer, 2024), given the number of transcripts affected and the directionalities of the differences between the mutants and the wild type (see 1 in Fig. 9G). Transcript levels of some other genes related to mineral nutrition, in particular nitrate (up), phosphate and sulfur (down), were additionally de-regulated in both *dph1-1* and *dph2-1*.

### *AtDPH1* and *AtDPH2* do not complement a yeast *dph1 dph2* double mutant

To test whether Arabidopsis *DPH1* and *DPH2* coding sequences can complement the *S. cerevisiae dph1* and *dph2* mutants, we made the constructs *ScDPH1*-(HA)_6_, *AtDPH1*-(HA)_6_, *ScDPH2*-(c-myc)_3_ and *AtDPH2*-(c-myc)_3_ (Fig. 10A). We then transformed the yeast *dph1* mutant with the *ScDph1*-(HA)_6_ or the *AtDph1*-(HA)_6_ construct, the yeast *dph2* mutant with the *ScDph2*-(c-myc)_3_ or the *AtDph2*-(c-myc)_3_ construct, and the yeast *dph1 dph2* double mutant with expression constructs for both *DPH1* and *DPH2*. Immunoblots detected all epitope-tagged proteins in the respective yeast genetic backgrounds (Supplementary Fig. S9A). Similar to Arabidopsis, yeast eEF2 protein is quantitatively diphthamide-modified (Hawer et al., 2018). In yeast *dph1*, *dph2*, and *dph1 dph2* double mutant cells, we detected unmodified eEF2 protein (Fig. 10B). We did not detect unmodified eEF2 in the yeast *dph1* mutant transformed with the *ScDPH1*-(HA)_6_ construct, in the yeast *dph2* mutant transformed with the *ScDPH2*-(c-myc)_3_, and in the yeast *dph1 dph2* double mutant transformed with both *ScDPH1* and *ScDPH2*, demonstrating the successful complementation of these mutants (Fig. 10B). However, the yeast *dph1* and *dph2* mutants expressing *AtDPH1*-(HA)_6_, *AtDPH2*-(c-myc)_3_, or both in combination, continued to accumulate unmodified eEF2 (Fig. 10B). Taken together, these results indicated that Arabidopsis DPH1-(HA)_6_ and DPH2(c-myc)_3_ were produced in yeast, but were unable to complement the *dph1* and *dph2* mutants.

**Figure 10.**
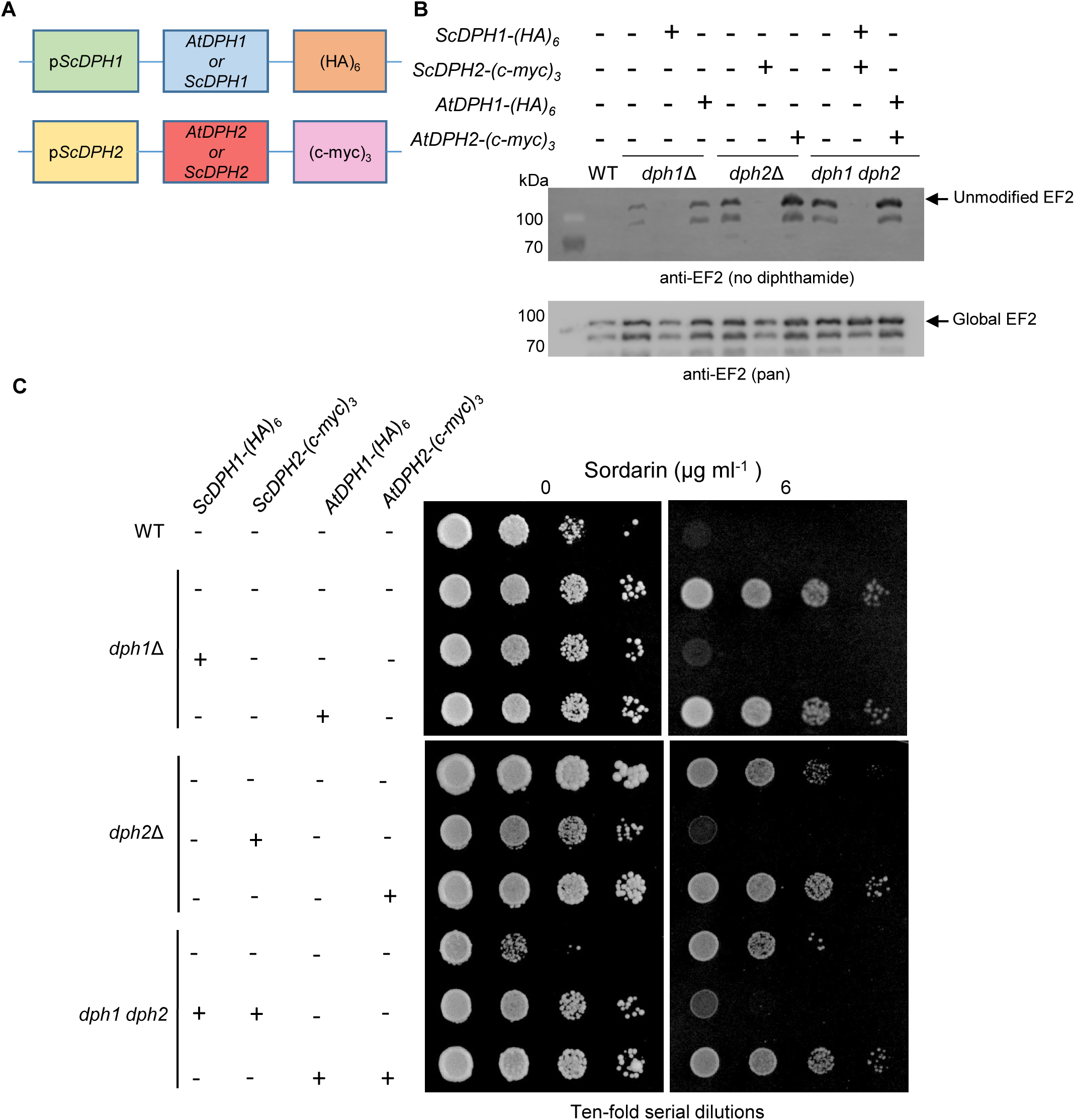
*A. thaliana DPH1* and *DPH2* do not complement a *S. cerevisiae dph1 dph2* mutant. **A)** Schematic overview of multicopy plasmid-based expression of *AtDPH1* and *AtDPH2* in yeast cells and immunoblot detection. The expression of *A. thaliana DPH1* and *DPH2* coding sequences was transcriptionally controlled by the promoter of the respective yeast homologue. *At*DPH1 was produced in yeast with a C-terminally fused to 6x HA-tag and *At*DPH2 with a C-terminal 3x c-myc tag. **B)** Immunoblot detecting cellular levels of total EF2 (global) and unmodified EF2 (no diphthamide). **C)** Phenotypic drop assay on media containing sordarin for the restoration of sordarin sensitivity upon *A. thaliana DPH* (co-)expression in the respective yeast deletion mutants. Yeast cells were transformed with either empty vectors or plasmids permitting the expression of *DPH* of either *A. thaliana* or *S. cerevisiae*. Transformants were spotted on YNB lacking leucine and uracil without (no toxin) or with addition of 6 µg ml^-1^ sordarin, and incubated at 30°C for 2 d.

The tetracyclic diterpene glycoside sordarin is an eEF2 inhibitor with antifungal properties (Domínguez et al., 1998). Sordarin targets eEF2 in a diphthamide-dependent manner to inhibit translation by blocking the eEF2-mediated translocation of tRNAs (Capa et al., 1998; Domínguez and Martín, 1998; Justice et al., 1998; Domínguez et al., 1999; Botet et al., 2008). Accordingly, wild-type yeast cells were sensitive to sordarin, while *dph1* and *dph2* yeast cells, both of which lack diphthamide, were tolerant to sordarin (Fig. 10C).

Sensitivity to sordarin was restored in the yeast *dph1* mutant transformed with the *ScDPH1*-(HA)_6_ construct, the yeast *dph2* mutant transformed with *ScDPH2*-(c-myc)_3_ construct and the yeast *dph1 dph2* double mutant transformed with both the *ScDPH1*-(HA)_6_ and the *ScDPH2*-(c-myc)_3_ constructs. However, yeast mutants remained tolerant to sordarin when transformed with the corresponding *AtDPH1* and *AtDPH2* constructs (Fig. 10C). This indicated that *AtDPH1*-(HA)_6_ and AtDPH2-(c-myc)_3_ failed to restore the sordarin sensitivity of the corresponding yeast mutants.

The DPH1 and DPH2 proteins of both yeast and Arabidopsis function as heterodimers (Dong et al., 2014). The presence of both Arabidopsis DPH1 and DPH2 failed to complement the diphthamide biosynthesis defect of a yeast *dph1 dph2* double mutant (Fig. 10B,C), although Co-IP assays suggested that AtDPH1-(HA)_6_ can interact with AtDPH2-(c-myc)_3_ in yeast cells, and that ScDPH1-(HA)_6_ interacts with ScDPH2-(c-myc)_3_ as a positive control (Supplementary Fig. S9B). Taken together, these results suggest that AtDPH1, AtDPH2, and also their combination, cannot functionally replace their respective orthologues in *Saccharomyces cerevisiae*.

## Discussion

### Functional relationship of DPH1 and DPH2 in Arabidopsis

The biosynthesis of diphthamide in archaea, yeast, mice, and humans begins with an unconventional radical SAM reaction (Liu et al., 2004; Zhang et al., 2010; Zhu et al., 2011; Thakur et al., 2012; Dong et al., 2014; Dong et al., 2017; Dong et al., 2018; Dong et al., 2019). The enzymes carrying out this reaction, the DPH2 homodimer in archaea and the DPH1-DPH2 heterodimer in eukaryotes, bind one 4Fe-4S cluster per subunit (Zhang et al., 2010; Zhu et al., 2011; Dong et al., 2017; Dong et al., 2018; Dong et al., 2019). Previously, we demonstrated the near-quantitative presence of the diphthamide modification on Arabidopsis eEF2 using mass spectrometry, showed that DPH1 is necessary for its presence, and we phenotypically characterized *dph1* mutants (Zhang et al., 2022). In this study, we report the conserved biochemical function of Arabidopsis DPH2 in plant diphthamide biosynthesis. We describe its conserved and its plant-specific molecular physiological functions, including the requirement of diphthamide for plant tolerance to oxidative stress and growth in the light, based on a reverse genetic approach.

In archaea, DPH2 forms a homodimer of two approximately V-shaped subunits interacting through the ends of both arms, with the clefts in each subunit harboring an 4Fe-4S cluster bound to three conserved cysteine residues, each of which is contributed by a different one of three distinct domains (Zhang et al., 2010). In this study, we modeled the structure of the AtDPH1-AtDPH2 complex, with a 4Fe-4S cluster in each subunit ligated by three cysteine residues. Only the positions of the first and the third cysteine residues are conserved in AtDPH2 by comparison to both the archaeal DPH2 and AtDPH1 (Fig. 1; Supplementary Figs. S1, S2). We further showed that AtDPH2 directly interacts with AtDPH1 by Co-IP and rBiFC (Fig. 2), suggesting the formation of a DPH1-DPH2 heterodimer in Arabidopsis, similar to the known dimeric complex in yeast (Supplementary Fig. S9B; Dong et al., 2014). The shared localization of *DPH1* and *DPH2* promoter activity and shared sub-cellular localization of the DPH1 and DPH2 proteins are consistent with their direct interaction (Fig. 4).

We were not able to complement yeast *dph1*, *dph2*, or *dph1 dph2* mutants by expressing the coding sequences of Arabidopsis *DPH1*, *DPH2* or *DPH1* together with *DPH2*, respectively (Fig. 10). DPH1 and DPH2 of both yeast and Arabidopsis must heterodimerize to form a functional radical SAM enzyme. There is a possibility that Arabidopsis AtDPH1 and AtDPH2 are not able to form heterodimers with yeast ScDPH2 and ScDPH1, respectively, given the overall low sequence similarity between the yeast and Arabidopsis orthologues (Supplementary Fig. S1). Moreover, it was reported that a total of five proteins, Dph1 to Dph4, together with Dph8, are involved in completing the first step of diphthamide biosynthesis in yeast (Arend et al., 2023; Schaffrath and Brinkmann, 2024). A recent study reported that the Arabidopsis AtDPH4 protein is an Hsp70 cochaperone with iron-binding activity, supporting a proposed function in diphthamide biosynthesis (Verma et al., 2024). Although we detected an interaction between Arabidopsis DPH1 and DPH2 also in yeast cells (Supplementary Fig. S9B), their heterodimeric complex might not be able to functionally interact with the remaining components of the multiprotein complex in the heterologous yeast expression system. Similar observations have occasionally been reported for subunits of other multi-protein complexes (Mehlgarten et al., 2010; Mehlgarten et al., 2017).

### Roles of diphthamide in the responses of Arabidopsis to environmental stress

Although diphthamide has been known for over 40 years (Van Ness et al., 1980), its biological role is only partly understood. So far, reports of growth defects or reproductive penalties in mutants lacking this unique post-translational modification were restricted to multicellular organisms such as mice (Chen and Behringer, 2004), human (Alazami et al., 2015; Loucks et al., 2015; Hawer et al., 2020), and recently Arabidopsis (Zhang et al., 2022), but not observed in the unicellular yeast (Hawer et al., 2018) under standard environmental conditions without further modifications. *S. cerevisiae dph* mutants showed increased sensitivity to heat stress (Hawer et al., 2018), and *dph^-/-^* mutants of the human MCF7 cell line accumulated elevated ROS levels even under normal growth conditions (Mayer et al., 2019). Different from yeast, we previously reported that Arabidopsis *dph1* mutants were more tolerant to heat stress, but they showed an activation of autophagy under standard growth conditions, and they were hypersensitive to heavy metal stresses from exposure to Cd and to excess Cu (Zhang et al., 2022). Therefore, it is tempting to speculate that part of the biological roles of diphthamide may be more easily detectable under certain stress conditions or specific cellular metabolic circumstances. A study on Chinese hamster ovary cells reported that eEF2/diphthamide functions in selectively translating internal ribosomal entry site-containing mRNAs that encode proteins necessary for cell survival under oxidative stress. Global eEF2 levels were significantly decreased in wild-type Chinese hamster ovary (CHO) cells under oxidative stress, and this decrease was more pronounced in diphthamide-deficient cells, indicating that diphthamide may play a role in protecting eEF2 against degradation under oxidative stress (Argüelles et al., 2014). Under conditions of high oxidative stress, diphthamide-deficient CHO cells were more sensitive to cell death (Argüelles et al., 2014). Similarly, eEF2 protein levels and activity are highly sensitive to oxidative stress in male Wistar rats (Argüelles et al., 2006; Argüelles et al., 2009; Argüelles et al., 2011). In the present study, we demonstrated that diphthamide on eEF2 plays a role in maintaining growth and survival of Arabidopsis under oxidative stress and under conditions of elevated light intensities (Figs. 6-8; Supplementary Figs. S7,S8), suggesting the possibility of a conserved function of diphthamide in cellular tolerance to oxidative stress. Moreover, diphthamide-unmodified eEF2 levels increased and global eEF2 protein levels decreased under oxidative stress also in Arabidopsis (Fig. 8D). The mechanisms leading to reduced TOR activity in *dph1* and *dph2* mutants remain to be elucidated (Fig. 5 and Supplementary Fig. S6).

We previously reported that Cu and Cd stress led to the accumulation of unmodified eEF2 protein in wild-type plants (Zhang et al., 2022). Besides the possibility that the strong ligand-binding affinities of intracellular heavy metal cations could affect the occupancy of sensitive Fe-S cluster binding sites including those of radical SAM enzymes, ROS might also decrease the Fe-S cluster occupancy of the AtDPH1-AtDPH2 complex, thus interfering with its function and diphthamide modification of eEF2 *in planta* (Krämer, 2024). We investigated the effect of long-term mild oxidative stress on diphthamide biosynthesis by growing wild-type seedlings on 5 nM MV for 18 d and observed a weak band corresponding to unmodified eEF2 (Zhang et al., 2022). In this study, we exposed wild-type seedlings to 12 µM MV for 2 d, which generated a burst of oxidative stress. Apparently, this treatment had a stronger effect on diphthamide biosynthesis (Fig. 8A). Exposure to heavy metal excess is well known to cause oxidative stress in plants. Therefore, we cannot exclude that the interference of Cu and Cd with the diphthamide modification of eEF2 is attributable to secondary oxidative stress rather than the primary displacement of the Fe-S cluster of AtDPH1 and AtDPH2 by heavy metal cations. Besides heavy metal stress, numerous other abiotic and biotic stresses can cause oxidative stress in plants. Recent evidence further underpins the pronounced oxygen sensitivity of DPH1 and DPH2 enzymes from yeast and human (Zhang et al., 2021; Baik et al., 2023). Under aerobic conditions, the 4Fe-4S cluster of ScDph1 undergoes oxidation to a 3Fe-4S cluster, and ScDph3 acts to restore a functional ScDph1 (Zhang et al., 2021). It is thus possible that various stresses affect plant growth and development through decreased diphthamide biosynthesis, which acts to relay an integrative cellular stress signal consisting of the oxidation of a sensitive 4Fe-4S cluster in the AtDph1-AtDph2 heterodimer. Our results indicate not only that cellular diphthamide biosynthesis can be decreased under conditions of oxidative stress (Fig. 8) and heavy metal excess (Zhang et al., 2022), but also that a lack of diphthamide sensitizes plants to oxidative stress and elevated light, as observed here in *dph1* and *dph2* single and double mutants (Figs. 6,7; Supplementary Fig. S7). Taken together, these processes have the potential to re-inforce one another in a feed-forward loop to promote cell death.

Besides its specific functions under stress conditions, diphthamide may also function as a site for the regulatory modification of eEF2. Mono-ADP-ribosyltransferases (mADP-RTs) constitute an enzyme family that cleave NAD^+^ and covalently attach an ADP-ribosyl moiety to substrate proteins (Wirthmueller and Banfield, 2012). Enzymatic mono-ADP ribosylation is an ancient mechanism to regulate protein function (Krueger and Barbieri, 1995; Pallen et al., 2001; Corda and Di Girolamo, 2003). In addition, mADP-RTs are important virulence factors of bacteria that infect mammals and can use diphthamide as a target. It has been postulated that ADP ribosylation by diphtheria toxin may reflect an endogenous cellular control mechanism that also operates in uninfected cells (Collier, 1975). In mammalian cells, an endogenous ADP-ribosyltransferase activity specific for eEF2 was reported, and it may control protein synthesis (Lee and Iglewski, 1984; Sitikov et al., 1984; Sayhan et al., 1986). Also in plant pathogenic bacteria, mADP-RTs were identified as important virulence factors (Wirthmueller and Banfield, 2012). Whether or not these bacteria target diphthamide during plant infection remains to be investigated.

### Towards molecular mechanisms mediating the effects of diphthamide in plants

The analysis of *dph1* and *dph2* mutants informs us on the alterations to expect under environmental conditions that decrease the proportion of diphthamide-modified eEF2, for example heavy metal exposure and oxidative stress (Zhang et al., 2022). Our data unequivocally support a role of diphthamide in translational fidelity also in Arabidopsis (Fig. 5c, Supplementary Fig. S6D), as demonstrated previously in yeast and mammals (Ortiz et al., 2006; Liu et al., 2012; Hawer et al., 2018; Zhang et al., 2022). Based on these results, we developed a set of hypotheses for transcriptome alterations in *dph1* and *dph2* mutants (Fig. 9G). For these considerations, it should be noted that the normalization to Renilla luciferase activity in our reporter assay masked any possible effects of NMD on the abundance of the dual reporter transcript, a technical aspect that precludes any quantitative predictions (see results section and below). Transcriptomics of *dph1* and *dph2* mutants revealed the deregulation of the abundances of as many as 2,800 transcripts, among which about 1,186 transcripts were shared between both mutants. In the future, studying additional allelic mutants under standardized conditions will help to identify whether there are any differences between the biological functions of *DPH1* and *DPH2*. De-regulated transcripts shared between *dph1* and *dph2* mutants were in general agreement with the phenotypes of the mutants. Overall, the levels of transcripts related to abiotic stress responses, leaf senescence and programmed cell death were elevated whereas the abundance of transcripts related to biotic stress responses and growth-promoting hormones were lower in the mutants than in the wild type. The levels of the iron deficiency responsive transcripts encoding transcription factors bHLH038 and bHLH101, which are known to increase also in response to salicylic acid, were consistent with this (Kang et al., 2003). Yet, the de-regulation of a broader set of Fe homeostasis-related transcripts rather suggested that the physiological Fe status of the mutants was more sufficient than that of the wild type. Transcriptional alterations for 78 genes (34 up-regulated in mutants, 44 down-regulated in mutants) were consistent with those previously reported in leaves of both *dph1-1* and *dph1-2* (Supplemental Data Set 2) (Zhang et al., 2022). We consider these as a core set of 78 robustly responding transcripts in *dph1* and *dph2* mutants, given that we observed these responses independently of plant age and cultivation system.

Our transcriptome analyses of *dph1* and *dph2* mutants provide some initial support for a role of NMD, which will need to be addressed, including the question of whether diphthamide contributes to counteracting transposon activity and the expression of natural antisense transcripts. In yeast, at least part of the cell biological and metabolic phenotypes resulting from a lack of diphthamide involve enhanced rates of ribosomal - 1 frameshifting error during the translation of specific transcripts carrying programmed frameshifting sites. Programmed frameshifting sites are well-characterized in yeast (Harger and Dinman, 2003; Zhang et al., 2021), but they remain poorly known and poorly predictable in plants. Ongoing advancements of ribosome footprinting techniques, among other approaches, may soon allow sufficient precision to address ribosomal frameshifting in *dph* mutants in a gene-specific manner.

In conclusion, here we explored the roles of Arabidopsis DPH2, which can physically interact with AtDPH1 and is required for diphthamide biosynthesis, likely by participating in the catalysis of the first committed step. We identified two *dph2* mutant alleles and report their phenotypes under normal growth conditions, under oxidative stress and in elevated light. Arabidopsis *dph2* mutants exhibited growth retardation, hypersensitivity to hygromycin, lack of diphthamide and elevated -1 frameshifting errors, indistinguishable from *dph1* single and *dph1 dph2* double mutants. These findings demonstrate the biological importance of AtDPH2 in maintaining normal growth rates and responses of Arabidopsis to abiotic environmental stress. The broad conservation of diphthamide biosynthesis, together with the defects in Arabidopsis *dph1* and *dph2* mutants, suggest that the diphthamide modification of eEF2 acts in the protection of cells from environmentally induced damage and in relaying physiological equivalents of environmental cues to plant metabolism and growth.

## Methods

### Plant materials and growth conditions

*Arabidopsis* Columbia-0 (Col-0, wild type), *dph1-1* (SALK_205272C), *dph2-1* (SALK_126982) and *dph2-2* (SALK_024814) in the Col-0 background were obtained from NASC (http://arabidopsis.info/). T-DNA positions were obtained from TAIR (https://www.arabidopsis.org/index.jsp). The *dph1 dph2* double mutant was generated by crossing *dph1-1* with *dph2-1*. A homozygous line was identified in the F2 by PCR-based genotyping and propogated to F3 for the experiments conducted here. For cultivation, seeds were surface-sterilized in chlorine gas for 3 h and sown on 120-mm square polystyrene petri dishes (Greiner Bio-One, Frickenhausen, DE) containing 0.5x MS salts (Murashige and Skoog, Duchefa, Haarlem, NL), with 1% (w/v) sucrose and 0.8% (w/v) agar (Type M, Sigma, Steinheim, DE) (addressed as 0.5x MS medium below). Plates were kept in the dark at 4°C for 2 d before transfer into a growth chamber with 12-h light (120 µmol photons m^-2^ s^-1^, 22°C) and 12-h dark (18°C; CLF Plant Climatics, Emersacker, Germany) in vertical orientation, unless indicated otherwise. For growing plants in soil, seeds were first germinated on agar-solidified 0.5x MS medium for 2 w, as described above before transfer into soil (Minitray, Balster Einheitserdewerk, Fröndenberg, DE) and a growth chamber at 12-h light and 12-h dark (temperatures and light intensity were the same as described above). Plants were grown in long days (16 h) for RT-PCR and the GUS staining experiments. For hygromycin sensitivity assays, sterilized seeds were sown on agar-solidified 0.5x MS medium with or without 2 µM hygromycin B (Duchefa) in round petri dishes (Greiner Bio-One) positioned horizontally. For methyl viologen (Sigma) treatment, sterilized seeds were germinated and cultivated on agar-solidified 0.5x MS medium for 11 d or 12 d before the transfer of seedlings to fresh 0.5x MS medium or 0.5x MS supplemented with various concentrations of methyl viologen for 2 or 3 d as indicated. For the elevated-light sensitivity assays on plates, 7-d-old seedlings grown on 0.5x MS medium under control conditions (16 h light/8 h dark, 100 µmol photons m^-2^ s^-1^) were transferred to elevated-light (16 h light/8 h dark, 145 µmol photons m^-2^ s^-1^). For light sensitivity assays in soil, 24-d-old plants grown on soil under control conditions (16 h light/8 h dark, 120 µmol photons m^-2^ s^-1^) were transferred into elevated light intensity (constant light, 185 µmol photons m^-2^ s^-1^). *Nicotiana benthamiana* were grown in a growth chamber at 22°C with 16 h light/8 h dark. Five to six-week-old plants were used for infiltration. All PCR primer sequences used in this study are listed in Supplementary Table 1.

### Sequence alignment, phylogenetic analysis, and modeling of the AtDPH1-AtDPH2 structure

The amino acid sequence of the DPH2 protein of *S. cerevisiae* served as a query sequence to retrieve DPH2 homologs from the NCBI protein database (https://www.ncbi.nlm.nih.gov/) using blastp (Altschul et al., 1990). Multiple sequence alignment and the following construction of a neighbor-joining tree of DPH2 homologs were performed with MEGA6.0 software (Tamura et al., 2013). Bootstrap analysis was conducted with 1,000 iterations. Structure modelling of AtDPH1-AtDPH2 complex was performed using the crystal structure of DPH2 from Archaea species *Pyrococcus horikoshii* (PDB/3LZD) as a template with online tool SWISS-MODEL (https://swissmodel.expasy.org/). The predicted model with the highest GMQE (Global Model Quality Estimate) score was selected (Waterhouse et al., 2018). The predicted structure was further processed with PyMOL including its output (Schrodinger LLC, 2015).

### Ratiometric bimolecular fluorescence complementation (rBiFC)

*AtDPH1* and *AtDPH2* coding sequences were cloned from cDNA by PCR into a pENTR/D-TOPO entry vector by BP reaction (Fisher Scientific). Subsequently, *AtDPH1* and *AtDPH2* coding sequences were transferred from entry vectors to a gateway-compatible binary 2in1 rBiFC vector (Grefen and Blatt, 2012) through the LR reaction (Fisher Scientific), and after sequence verification the resulting constructs were used in transient transfection of *N. benthamiana* leaf epidermal cells employing syringe-mediated agro-infiltration as described previously (Mehlhorn et al., 2018). Fluorescence intensities were quantified at 3 d post-infiltration with a Leica SP5 confocal laser scanning microscope (YFP at 514 nm excitation (ex) and 520–560 nm emission (em); RFP at 561 nm excitation and 565–620 nm emission). YFP/RFP ratios were calculated from 25 different regions of leaves.

### Co-immunoprecipitation (Co-IP) analysis

*AtDPH1* and *AtDPH2* coding sequences were transferred from entry vectors (see above) to a gateway-compatible FRET 2in1 destination vector containing monomeric enhanced green fluorescent protein (mEGFP) and mCherry (pFRETgc-2in1) (Hecker et al., 2015; Mehlhorn et al., 2018) through the LR reaction (Fisher Scientific) to generate the constructs *p35S:DPH1-mCherry-p35S:DPH2-eGFP* and *p35S:DPH1-mCherry-p35S:DPH1-eGFP* (Fig. 2a). After verification by sequencing, the resulting constructs were used for transient transfection of *N. benthamiana* leaf epidermal cells for subsequent Co-immunoprecipitation (Co-IP) of fusion proteins (Hecker et al., 2015; Mehlhorn et al., 2018). Around 200 mg leaf material was harvested followed by flash freezing at 3 d post-infiltration and homogenized in liquid nitrogen. Then 400 μL lysis buffer (25 mM Tris pH 8.0, 150 mM NaCl, 1% (v/v) Nonidet P-40, and 0.5% (w/v) Na-deoxycholate) supplemented with 1× Pierce protease inhibitor (Fisher Scientific) and 2 mM DTT was added, followed by incubation at 4°C for 1 h. After centrifugation, the supernatant was mixed with 20 to 25 μL of RFP beads (RFP-trap; Chromotek) and then incubated at 4°C for 1 h. Beads were collected by centrifugation, transferred onto spin columns, and rinsed twice with lysis buffer, followed by six washes with wash buffer (25 mM Tris pH 8.0, 150 mM NaCl). Co-immunoprecipitated proteins were eluted with 2× Laemmli buffer containing 3.5% (v/v) β-mercaptoethanol and heated at 95°C for 5 min. Proteins were separated by 10% (v/v) SDS-PAGE and detected by Western blots (see Immunoblots section).

### RT-PCR and RT-qPCR

Total mRNA was extracted using the RNeasy Plant Mini Kit (Qiagen) kit following the manufacturer’s protocol. Reverse transcription was carried out on 1 μg total RNA after DNase I-digestion (New England Biolabs) using the SuperScript III kit (Fisher Scientific) and oligo(dT) primers following manufacturer’s instructions. For RT-PCR, first-strand cDNAs were used as templates for PCR using DreamTaq Green DNA-Polymerase (Fisher Scientific) and detected by agarose gel electrophoresis. RT-qPCR was performed with first-strand cDNAs using GoTaq qPCR Master Mix (Promega) with a LightCycler480 (Roche). Transcript levels were calculated with the 2^-ΔCt^ method and normalized to *UBQ10* expression levels. Primer sets used for RT-PCR and RT-qPCR are given in Supplementary Table 1.

### Plasmid construction for stable *A. thaliana* transformation

For genetic complementation of the Arabidopsis *dph2-1* mutant and DPH2 subcellular localization, the construct *pDPH2:DPH2-GFP* was generated with GreenGate cloning (Lampropoulos et al., 2013). Briefly, the promoter region amplified by PCR with Arabidopsis genomic DNA as template (1.8 kb upstream of the translational start codon), the coding sequence, and the 3’ UTR sequence (281 bp downstream of the stop codon) of *DPH2* amplified by PCR with Arabidopsis cDNA as template were separately cloned into the appropriate pUC19-based entry vectors *via BsaI* sites. The PCR was performed using the Phusion High-Fidelity DNA Polymerase (Fisher Scientific) following the manufacturer protocol with a Thermal Cyclers (Bio-Rad). Subsequently, the three completed entry vectors, together with pGGB003 (*B-dummy*), pGGD001 (*linker-eGFP*), and pGGF008 (*pNOS:BastaR:tNOS*), were assembled into a pGreen-IIS based destination vector PGGZ003 via the *BsaI* sites to obtain a translational *eGFP* fusion construct of *DPH2* under the control of the native promoter. The two completed entry vectors containing the 1.8-kb promoter region and the 281-bp 3’ UTR region of DPH2, pGGB003 (*B-dummy*), pGGC051 (*GUS*), pGGD002 (*D-dummy*), and pGGF008 (*pNOS:BastaR:tNOS*), were assembled into the pGreen-IIS based destination vector PGGZ003 to generate the *pDPH2:GUS* construct employing GreenGate cloning (Lampropoulos et al., 2013), followed by the stable transformation of wild-type Arabidopsis (Col-0) plants. *Agrobacterium tumefaciens*-mediated transformation using the strain GV3101 was conducted as described (Clough and Bent, 1998).

### Immunoblots

Tissue powder (100 mg) was homogenized in 200 µl of Laemmli buffer (125 mM TRIS-HCl pH 6.8, 10% (v/v) mercaptoethanol, 0.01% (v/v) bromophenol blue, 4% (w/v) SDS, and 20% (v/v) glycerol) and incubated at 95°C for 10 minutes. After centrifugation, supernatants (10 µl of protein extract per lane) were separated by 10% (w/v) SDS-PAGE and blotted onto a polyvinylidene difluoride (PVDF) membrane by wet/tank transfer. For the detection of global and diphthamide-unmodified eEF2, the PVDF membrane was blocked with 5% (w/v) BSA (bovine serum albumin) in TBS-T (0.05% Tween-20). The blocked membrane was probed with global eEF2 antibody (3C2, 1:8,000, Roche) or diphthamide-unmodified eEF2 antibody (10G8, 1:8,000, Roche), followed by incubation with the secondary antibody goat anti-rabbit IgG/HRP (1:10,000, 12-348, Fisher Scientific). For the detection of Arabidopsis ACTIN, GFP/DPH1-GFP and GFP/DPH2-GFP, the PVDF membrane was blocked with 5% (w/v) low-fat milk powder (AppliChem) in TBS-T (0.1% Tween-20). The blocked membrane was probed with anti-ACTIN (1:8,000, AS13 2640, Agrisera, Vännäs, SE) or anti-GFP (1:5,000, AB10145, Sigma), followed by incubation in the secondary antibody goat anti-rabbit IgG/HRP (Fisher Scientific). For the immunoblots in Co-IP, the PVDF membrane was blocked with 5% (w/v) low-fat milk powder (AppliChem) in TBS-T (0.1% Tween-20). The blocked membrane was probed with anti-RFP (1:2,500, Chromotek), anti-GFP IgG1κ (1:1,000, Roche), followed by incubation in the secondary antibody goat anti-mouse IgG [Fc-specific] (1:10,000, Sigma-Aldrich). Signals were detected with the ECL Select Western Blotting Detection Reagent (GE Healthcare, Little Chalfont, UK) and a Fusion Fx7 GelDoc (Vilber Lourmat, Eberhardzell, DE). All membranes were blocked at room temperature for 1 h, and all antibodies were added to a solution of the same composition as used for blocking followed by incubation at RT for 1 h.

### DPH2 subcellular localization

Roots of 9-d-old seedlings of *dph2-1 pDPH2:DPH2-GFP* complementation lines were stained with 0.01 g l^-1^ propidium iodide (Sigma) for 1 min and briefly washed in ultrapure water twice. GFP florescence was detected using a Leica SP5 confocal laser scanning microscope, with an excitation wavelength at 488 nm and detection using GFP settings (GFP at 488 nm excitation and 500–536 nm emission). Three T3 seedlings were imaged for each of the three independently transformed complementation lines. Wild type (Col-0) was used as a negative control. To generate the construct *pUBQ10:DPH2-GFP-T35S* for subcellular localization of DPH2-GFP fusion in protoplasts, the *AtDPH2* coding sequence amplified from cDNA was cloned into a pENTR/D-TOPO entry vector through the BP reaction (Fisher Scientific). Subsequently, the *DPH2* coding sequence was transferred from the entry vector to the gateway-compatible vector pUBC-GFP-Dest (Grefen et al., 2010) through the LR reaction (Fisher Scientific). The construct *pUBQ10:DPH2-GFP-T35S* was transfected into Arabidopsis mesophyll protoplasts isolated from 4-w-o soil-grown wild-type plants for transient expression assay as described (Yoo et al., 2007). DPH2-GFP fusion was expressed in protoplasts by incubating at 22°C under constant light for 16 h. The transfected and untransfected protoplasts (control) were observed with a Leica SP5 confocal laser scanning microscope, with an excitation wavelength at 488 nm and detection using GFP settings for GFP fluorescence (Leica, Wetzlar, DE) or propidium iodide settings for chlorophyll autofluorescence.

### Frameshifting assays in protoplasts

Frameshifting assays were performed exactly as described (Zhang et al., 2022). A dual-luciferase reporter system developed for yeast with a control reporter (pJD375) and a LA-Virus site-based -1 frameshift reporter (pJD376) (Harger and Dinman, 2003) was adapted for transient expression assays in Arabidopsis mesophyll protoplasts. The *renilla* and *firefly luciferase* cDNAs, together with either the polylinker region from pJD375 or the programmed frameshifting signal from pJD376, were amplified with primers introducing *XhoI* and *SpeI* sites. The amplified fragments were cloned into the *XhoI* and *SpeI* sites of transient expression vector pMatrix (Grefen et al., 2010). The resulting plasmids were used to transfect mesophyll protoplasts isolated from soil-grown wild-type, *dph2*, and *dph1 dph2* mutant plants as described (Yoo et al., 2007). Transfected protoplasts were harvested by centrifugation after dark incubation at 22°C for 16 h. Protoplasts were disrupted through two freeze-thaw cycles. Firefly and renilla luciferase activities were quantified using the dual-luciferase reporter assay system (Promega) in a Synergy HTX microplate reader (BioTek, Bad Friedrichshall, DE).

### RNA-sequencing and data analysis

RNA extraction, sequencing, and data analyses were performed as described previously (Zhang et al., 2022). Five two-week-old seedlings cultivated in 2 ml liquid 0.5x MS salts with 0.5% (w/v) sucrose in a six-well plate (Greiner Bio-One) with shaking (100 rpm min^-1^) under 12-h light (120 µmol photons m^-2^ s^-1^, 22°C) and 12-h dark (18°C; CLF Plant Climatics, Emersacker, Germany) cycles were harvested per sample approximately at ZT 6 for total RNA isolation. Libraries were prepared and sequenced using the Illumina platform by Novogene (Cambridge, UK). Raw reads were first filtered by CutAdapt 2.1 to remove adaptors and low-quality reads. The filtered reads were mapped to the *Arabidopsis thaliana*Col-0 reference genome TAIR10 (https://www.arabidopsis.org/download/list?dir=Sequences%2FTAIR10_blastsets). Read counts were determined by Qualimap2 (Okonechnikov et al., 2016). Differentially expressed genes were identified using the R package *DESeq2* (Love et al., 2014). AgriGO (http://systemsbiology.cau.edu.cn/agriGOv2/index.php) was used for gene ontology (GO) term enrichment analyses with the Yekutieli (FDR under dependency) multi-test adjustment method (Tian et al., 2017).

### Detection of GUS activity by histochemical staining

Freshly harvested tissues of homozygous T3 *pDPH2:GUS* lines were immersed in GUS staining buffer (0.2 % (v/v) Triton X-100, 50 mM sodium phosphate buffer pH 7.2, 2 mM potassium ferrocyanide, 2 mM cyclohexylammonium 5-bromo-4-chloro-3-indolyl-β-D-glucuronate, X-Gluc) at 37°C for 1 d after 5 min vacuum infiltration. The samples were then immersed in 75% (v/v) ethanol overnight to remove chlorophyll before imaging with a microscope (VS120, Olympus, Hamburg, DE).

### Chlorophyll measurement

Total chlorophyll was extracted from 20 mg frozen powder with 2 ml methanol and quantified spectrophotometrically at 652 nm and 665 nm with a Synergy HTX microplate reader (BioTek, Bad Friedrichshall, DE). Chlorophyll content was calculated using the equation: Chlorophyll (µg ml^-1^) = 22.12 x A_652 nm_ + 2.71 x A_665 nm_ (Porra, 1989).

### Yeast strains, plasmid generation and yeast-based assays

Yeast strains BY4741 WT, *dph1*Δ*::kanMX* and *dph2*Δ*::kanMX* were obtained from the Euroscarf KO collection. For the generation of *dph1*Δ*dph2*Δ*, DPH2* was deleted in the *dph1*Δ*::kanMX* background strain using a PCR-mediated *SpHIS5* gene deletion cassette derived from pUG27 (Gueldener, 2002). For multicopy plasmid-based expression of *ScDPH1* and *ScDPH2*, genomically epitope-tagged alleles (Janke et al., 2004) were amplified from yeast strains via PCR including native promoter and terminator regions of *ScDPH1* and *ScDPH2*. Subsequently, PCR products were cloned into multicopy plasmids YEplac181 and YEplac195 (Gietz and Sugino, 2015), respectively, using the optimal cloning method (Jacobus and Gross, 2015). For plasmid-based *AtDPH1* and *AtDPH2* expression in yeast, Arabidopsis cDNA fragments comprising the respective open-reading frame were cloned into the aforementioned plasmids to substitute *ScDPH1* and *ScDPH2* open reading frames respectively. Yeast strains were then transformed with plasmids using the lithium-acetate transformation method (Gietz and Woods, 2002) and selected on minimal yeast nitrogen base media lacking leucine and uracil as appropriate. Phenotypic assays were performed by spotting 10-fold dilutions of cell suspensions starting at an optical density (600 nm) of 1 on yeast nitrogen base media lacking leucine and uracil containing 6 μg μL^-1^ sordarin sodium salt (Sigma-Aldrich, St. Louis, Missouri, USA). Yeast protein extraction and co-immune precipitation were conducted as previously reported (Krutyholowa et al., 2019). Protein samples were separated by SDS-PAGE and analyzed by Western blotting using anti-HA (2-2.2.14, Invitrogen, Waltham, Massachusetts, USA), anti-c-myc (9E10, kindly provided by Prof. Dr. Markus Maniak, University of Kassel, Germany), anti-Cdc19 serum (kindly provided by Dr Jeremy Thorner, University of California, Berkley, CA, USA), anti-EF2(pan) and anti-EF2(no diphthamide) antibodies as described before (Hawer et al., 2018).

### Statistical analyses

All statistical analyses were done by IBM SPSS Statistics (Version 29.0.2.0). All data shown in graphs and tables are represented as the mean ± s.d., from at least 3 replicates. Comparisons of means were done by two-tailed Student’s *t*-test (*, *P* < 0.05, **, *P* < 0.01, ***, *P* < 0.001) or one-way ANOVA with Tukey’s test, Waller-Duncan, or Games-Howell test (*, *P* < 0.05, **, *P* < 0.01, ***, *P* < 0.001).

### Accession numbers

RNA-seq data are available from ArrayExpress (www.ebi.ac.uk/arrayexpress, E-MTAB-13779). Accession numbers are as follows: *AtDPH1* (AT5G62030.1, TAIR10), *AtDPH2* (AT3G59630.1, TAIR10), *AtDPH3* (AT2G15910), *AtDPH4* (AT4G10130), *AtDPH5* (AT4G31790), *AtDPH6* (AT3G04480), *AtDPH7* (AT5G63010), *AtDPH8* (AT1G27060), *eEF2* (AT1G56070), *TOR* (AT1G50030), *S6K* (AT3G08720), *ScDPH1* (YIL103W), *ScDPH2* (YKL191W).

## Supplementary data

**Supplementary Figure S1. Diphthamide biosynthesis pathway and identification of homologues in Arabidopsis. A**, Schematic representation of the diphthamide biosynthetic pathway, organized into steps I to III, showing the corresponding human and yeast proteins with their putative homologues in Arabidopsis and percentage identical amino acids according to sequence alignments. **B**, Multiple sequence alignment of DPH2 homologues. The three conserved Cys residues (Cys71, Cys92, Cys335) thought to be critical for 4Fe-4S cluster binding by eukaryotic DPH2 proteins are boxed in red. Red arrows mark amino acid positions in AtDPH2 that are at the predicted to localize to the interface between AtDPH2 and AtDPH1. Orange triangles mark amino acids possibly interacting with eEF2. The representative gene model AT3G59630.1 was used for sequence alignment. At, *Arabidopsis thaliana*; Os, *Oryza sativa*; Sm, *Selaginella moellendorffii*; Pp, *Physcomitrium* (*Physcomitrella*) *patens*; Hs, *Homo sapiens*; Sc, *Saccharomyces cerevisiae*. Poorly aligned C-terminal residues are not shown (At 111, Os 111, Sm 41, Pp 36, Hs 97, Sc 121 amino acids).

**Supplementary Figure S2. Sequence alignment of AtDPH1 and AtDPH2.** Vertical arrows mark amino acid positions in AtDPH1 (blue) and AtDPH2 (red) that are at the predicted to localize to the interface between AtDPH2 and AtDPH1. Cys residues predicted to bind the 4Fe-4S cluster are boxed in red (see Fig. 1). The residues contributing to the predicted SAM binding site of AtDPH1 are boxed in orange.

**Supplementary Figure S3. Identification of Arabidopsis *dph2* mutants and complementation of the *dph2-1* knock-out mutant. A**, Schematic representation of T-DNA insertions at the *DPH2* locus in *dph2-1* and *dph2-2* mutants. LB, T-DNA left border. Arrowheads point to the 5’ ends of the primers used for both RT-PCR (**B**) and RT-qPCR (**C**) (F: forward primer; R: reverse primer). **B**, Detection of *DPH2* transcripts (30 cycles) in leaf tissues of 4-week-old soil-grown wild-type, *dph2-1* and *dph2-2* plants by RT-PCR. *ACTIN8* was used as a constitutively expressed control gene (26 cycles). Each of the three lanes per genotype corresponds to plants cultivated in an independent experiment. **C**, RT-qPCR quantification of *DPH2* transcript levels in shoots of 4-w-old soil-grown wild-type, *dph2-1 and dph2-2* mutant lines, and three *dph2-1 pDPH2:DPH2-GFP* complemented lines. Shown are mean ± s.d. (*n* = 3 replicate pools generated from 5 plants per pool). Significant differences from WT: *, *P* < 0.05, ***, *P* < 0.001, one-way ANOVA with Tukey’s test. The no-template control gave a signal equivalent to the one observed for *dph2-1*. **D**, Immunoblot detection of the DPH2-GFP fusion protein in shoots of 4-w-old soil-grown wild-type and *dph2* mutants (negative controls), as well as *dph2-1 pDPH2:DPH2-GFP* complemented lines (*pDPH2*: *DPH2* promoter). Total protein extracts were resolved by SDS-PAGE, blotted and probed with an anti-GFP antibody. Same blot stained with Coomassie Brilliant Blue (CBB, bottom image) served as a loading control.

**Supplementary Figure S4. Smaller size of *dph1*, *dph2* and *dph1 dph2* double mutants in soil, and hygromycin sensitivity**. **A**, Representative photographs showing 31-d-old wild-type (WT), *dph1-1*, *dph2* mutants, the *dph1-1 dph2-1* double mutant, and *dph2-1* complemented lines, cultivated in soil. Scale bars, 2 cm. **B**, Photograph of two-week-old seedlings of WT, *dph1-1, dph2-1*, and *dph2-2* cultivated in agar-solidified 0.5x MS medium containing 1% (w/v) sucrose without (control) or with 2 μM hygromycin B (hyg).

**Supplementary Figure S5. AtDPH2 localization in the cytosol.** Representative confocal image of a protoplast transiently transfected with *pUBQ10:DPH2-GFP-T35S* (top row) as well as an untransfected protoplast (control, bottom row). Red color represents autofluorescence of chloroplasts. BF, bright field. Scale bars, 20 μm. The images shown are representative of 6 imaged protoplasts in total from 2 independent transfections.

Supplementary Figure S6. Reduced TOR activity in *dph1* and *dph2* mutants and enhanced rates of translational -1 frameshifting error. A, Ratios of firefly to renilla luciferase activities in Arabidopsis mesophyll protoplasts. Mesophyll protoplasts isolated from the wild type (WT), *dph2*, and *dph1 dph2* mutants were transfected with pYDL-L-A (control) and pYDL-L-A(-1) (test), respectively (see Fig. 5B). Data are mean ± s.d. (*n* = 3 independently transformed replicate aliquots of protoplasts). Significant differences from WT: *, *P* < 0.05, one-way ANOVA with Tukey’s test. **B**-**D**, Additional immunoblots used for calculating TOR activity according to the levels of S6K-P and S6K in 14-d-old wild-type (WT), *dph1*, and *dph2* mutant seedlings (see Fig. 5E). Total protein was extracted from equal amounts of biomass, resolved by SDS-PAGE, blotted and probed with antibodies against S6K-P and S6K, respectively. The positions of the S6K bands are indicated by red arrows. Each panel is from one independent experiment (**B**-**D**). Seedlings were cultivated in liquid 0.5x MS medium supplemented with 0.5% (w/v) sucrose. WT seedlings treated with 2 μM TOR inhibitor AZD-8055 (AZD) for 1 d were used as controls.

**Supplementary Figure S7. *Dph1* and *dph2* mutants are hypersensitive to elevated light.** Representative photographs of 30-d-old wild-type (WT), *dph1-1*, *dph2-1* and *dph1-1 dph2-1* double mutants grown in soil under control conditions (16 h light/8 h dark, 120 µmol photons m^-2^ s^-1^) and elevated-light conditions (constant light, 185 µmol photons m^-2^ s^-1^). Plants pre-cultivated under control conditions were either transferred to elevated-light conditions or further cultivated under the initial growth conditions for 6 d. Scale bars, 2 cm. Arrows indicate chlorotic leaves of *dph1-1*, *dph2-1*, and *dph1 dph2* mutants grown at an elevated light intensity.

**Supplementary Figure S8. Methyl viologen treatment reduces the levels of DPH1-GFP and DPH2-GFP fusion proteins, with *DPH1* and *DPH2* transcript levels unchanged. A,** Immunoblot detection of the DPH1-GFP fusion protein in the *dph1-1* complemented line Y1-5 (left) and of the DPH2-GFP fusion in the *dph2-1* complemented line Y1-3 (right) grown under control and MV stress conditions. Eleven-d-old seedlings of Y1-5 and Y1-3 grown on agar-solidified 0.5x MS medium with 1% (w/v) sucrose were transferred to the same medium without (control) or with 12 µM MV. Seedlings were harvested 2 and 3 d after transfer. Wild-type (WT), Y1-5, and Y1-3 seedlings harvested 3 d after transfer to fresh medium were used as controls. Total protein extracts were separated by SDS-PAGE, blotted and probed with an anti-GFP antibody. Numbers below the blot images indicate band intensities relative to those in Y1-5 or Y1-3 grown under the control condition, all normalized to the corresponding intensity of the actin band. **B**, RT-PCR of *DPH1* to *DPH7* transcript levels in WT seedlings grown under the conditions described above for 3 d. Samples from three independent cultivations (1 to 3) were used for RT-PCR. *DPH1* to *DPH7* (30 cycles), *ACTIN8* (26 cycles).

**Supplementary Figure S9. *At*DPH1 and *At*DPH2 interact in yeast cells. A**, Immunoblot showing relative protein amounts of *At*DPH1-(HA)_6_ in comparison to *Sc*DPH1-(HA)_6_ (anti-HA) as well as *At*DPH2-(c-myc)_3_ compared to *Sc*DPH2-(c-myc)_3_ (anti-c-myc). The yeast pyruvate kinase Cdc19 (anti-Cdc19) served as a loading control. **B**, Co-immunoprecipitation (Co-IP) assays of DPH1 and DPH2. A yeast *dph1 dph2* double mutant strain was (co-)transformed with plasmids constructed for the expression of epitope-tagged DPH1 and DPH2 proteins from either *A. thaliana* or *S. cerevisiae*. Total protein extracts were subjected to Co-IP using magnetic beads covalently coupled to anti-c-myc antibodies. IPs were examined by immunoblotting using anti-c-myc (to detect DPH2) and anti-HA antibodies (to detect co-immunoprecipitated DPH1)(left). A immunoblot of total protein extracts of the examined proteins prior to the IP (input) is shown on the right.

**Supplementary Table 1. Primers used in this study.**

**Supplemental Data Set 1. All expressed genes.**

**Supplemental Data Set 2. All differentially expressed genes.**

**Supplemental Data Set 3. Enriched GO terms.**

## Acknowledgements

We thank U. Brinkmann (Roche Innovation Center Munich, Penzberg, Germany) for the anti-eEF2 antibodies, and J. D. Dinman (University of Maryland, MD, USA) for sharing the plasmid for the -1 frameshifting assay, as well as Markus Maniak (University of Kassel, Germany) and Jeremy Thorner (University of California, Berkeley, USA) for sharing anti-c-Myc (9E10) and anti-Cdc19 antibodies, respectively.

In our research group, we thank H. Seebach for assistance with DPH1-DPH2 heterodimer modeling, and A. Aufermann and M. Pullack for technical assistance with plant growth.

## Author contributions

H.Z., and U.K. designed the research. H.Z. performed most experiments. N.J. analyzed the RNA-seq data. Yeast experiments were done by K.Ü. and T.M., L.Z. contributed to the protein interaction assays, L.W. contributed to the chlorophyll content assay and the RNA-seq experiment. H.Z., C.G., R.S., and U.K. analyzed data. H.Z. and U.K. wrote the manuscript. All authors edited the manuscript.

## Compliance and ethics

The authors declare that they have no competing interests.

